# Active Anemosensing Hypothesis: How Flying Insects Could Estimate Ambient Wind Direction Through Sensory Integration & Active Movement

**DOI:** 10.1101/2022.03.31.486300

**Authors:** Floris van Breugel, Renan Jewell, Jaleesa Houle

## Abstract

Estimating the direction of ambient fluid flow is a crucial step during chemical plume tracking for flying and swimming animals. How animals accomplish this remains an open area of investigation. Recent calcium imaging with tethered flying *Drosophila* has shown that flies encode the angular direction of multiple sensory modalities in their central complex: orientation, apparent wind (or airspeed) direction, and direction of motion. Here we describe a general framework for how these three sensory modalities can be integrated over time to provide a continuous estimate of ambient wind direction. After validating our framework using a flying drone, we use simulations to show that ambient wind direction can be most accurately estimated with trajectories characterized by frequent, large magnitude turns. Furthermore, sensory measurements and estimates of their derivatives must be integrated over a period of time that incorporates at least one of these turns. Finally, we discuss approaches that insects might use to simplify the required computations, and present a list of testable predictions. Together, our results suggest that ambient flow estimation may be an important driver underlying the zigzagging maneuvers characteristic of plume tracking animals’ trajectories.

## I. Introduction

Estimating the direction of ambient flow is a critical step for both swimming and flying animals during chemical plume tracking behaviors [1]. Without access to stationary flow sensors, however, the ambient direction of fluid flow cannot be directly measured [2]. How animals resolve this challenge remains an open area of investigation. Recent experimental work with fish has shown that they use temporal changes of flow from mechanosensory measurements to deduce flow direction during rheotaxis [3]. Comparatively less is known about how insects resolve this challenge while flying in windy environments.

Flying insects can measure *apparent wind*, corresponding to the vector sum of the ambient wind and motion induced wind. Decoupling apparent wind into its component parts is a straightforward vector subtraction if given absolute trajectory information in the same units as apparent wind. Insects, however, do not have direct access to such information. It is conceivable for an insect to estimate absolute ground speed using optic flow and inertial measurements (e.g. [4]), or to estimate ambient wind direction using a combination of optic flow and apparent wind [5]. However, both such approaches rely on calibrated measurements of the magnitude of optic flow and other modalities. Optic flow responses of insects, however, have depth- [6], texture- [7], temperature- [8], and state- [9] dependent response properties, resulting in constantly varying and unknown sensor gains. The magnitude of other sensor modalities will have similar time-dependent properties.

Through calcium imaging experiments, it has been found that a central part of insects’ brains, the central complex, encodes 2-dimensional angular compass information such as head orientation from visual landmarks [10] or polarization [11], apparent wind angle from their antennae [12], [13], and angular direction of motion from vision [14]–[16]. Neuroanatomical studies suggest that similar sensory encodings exist in many invertebrates [17]. These 2D angular encodings do not suffer from the same calibration challenges as magnitude measurements would, since the angular directions are equivalent to calculating the ratio of two perpendicular magnitude measurements, eliminating the unknown sensor gain. This feature and the physical overlap in the brain of the aforementioned sensory modalities presents an attractive hypothesis that perhaps they play a central role in insects’ ability to estimate ambient wind direction.

One strategy that relies solely on such angular measurements is called visual anemotaxis [18]. This algorithm would allow insects to orient upwind by turning until the angle of perceived apparent wind is aligned with the angle of motion. Behavioral experiments with free and tethered flies, however, suggest that their turns are often open-loop and ballistic in nature, and not actively controlled throughout the turn itself [19]–[21]. Furthermore, their free flight plume tracking behavior indicates that they are capable of moving crosswind at arbitrary body orientations with respect to the wind direction [22]. Together, these behavioral experiments suggest that insects may be capable of estimating the direction of ambient wind throughout their flight trajectories.

A recent nonlinear observability analysis of the three key sensor modalities encoded in the central brain of flies–orientation, apparent wind angle, and direction of motion–found that this sensor suite is mathematically sufficient for estimating the direction of ambient wind without explicitly requiring upwind orientation [23]. Estimating ambient wind direction in this manner, however, is only feasible for certain trajectories, such as those where the insect changes course direction.

This manuscript provides a detailed mathematical and simulation analysis to determine which trajectories offer the most accurate ambient wind direction estimates, as well as which computations are actually required to make these estimates. First we establish the mathematical relationships required to estimate ambient wind direction that are applicable to any estimator using the three measurements we consider. Second, we describe an optimal framework for estimating the ambient wind direction and validate the robustness of our approach using data from a flying drone. Third, we use our optimal framework to determine which types of trajectories are capable of providing the most accurate estimates under a variety of conditions. By using an optimal framework to answer this question we ensure that our results are equally applicable to other computationally simpler estimation algorithms designed for the same task. Finally, we discuss how insects might simplify the computations required to estimate ambient wind direction.

## II. Mathematics of Estimating Ambient Wind Direction

Inspired by the 2-dimensional angular encodings found in *Drosophila*, our analysis will focus on the plane parallel to the ground. Since this is also the plane where the largest changes in wind direction are usually observed [24], it also represents the most important angle that insects might need to estimate. Additional considerations for extensions to 3D are given in [23].

In this section we first review visual anemotaxis, and then develop a general framework for understanding how ambient wind direction *ζ*(*t*) could be estimated. In both cases we restrict our analysis to only consider the kinematic relationships and time history of three angular measurements (Fig. 1),

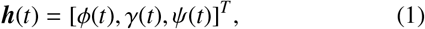

where *ϕ*(*t*) is the head orientation with respect to a global coordinate frame, *γ*(*t*) is the angular direction of airspeed relative to the head coordinate frame (equivalent to apparent wind +180°), and *ψ*(*t*) is the angular direction of motion relative to the head coordinate frame. For notational simplicity we drop the (*t*) going forward.

**Fig. 1:**
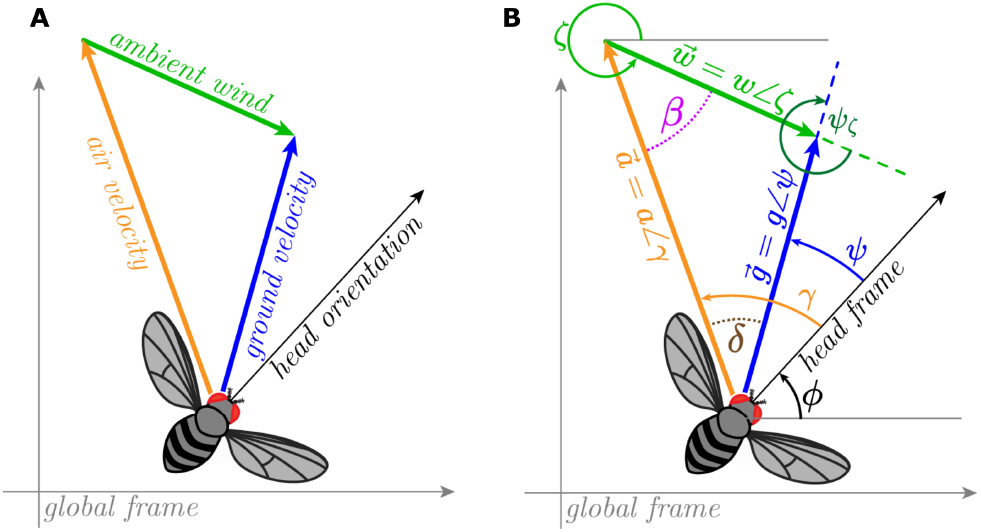
Trigonometric relationships of the angular sensory measurements.

We begin by defining the relationship between the angular measurements ***h***, the groundspeed (*g*), and wind speed (*w*) provided by the Law of sines (see Fig. 1):

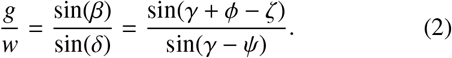

This relationship highlights the challenge of using the airspeed direction *γ* to estimate the wind direction, as any changes in the value of *γ* could be the result of changes in *ζ, g*, or *w*, none of which are available to the insect.

Since we are only concerned with estimating *ζ*, we simplify the relationship by defining *v* = *g/w*, a non-dimensional velocity describing the ratio of ground speed and wind speed, yielding:

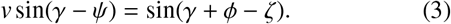

### A. Visual Anemotaxis (a review)

Before defining our framework for active anemosensing, we review the mathematics of the visual anemotaxis hypothesis. Consider what would happen if a fly were to actively control its orientation (*ϕ*) and direction of motion (*ψ*) based on sensory feedback:

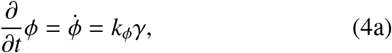

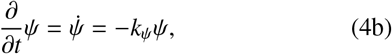

where *k_ϕ_* and *k_ψ_* are both constant gains that determine the rate of change in *ϕ* and *ψ*. Together, these control laws will ensure that the fly is oriented in the direction of its airspeed (*γ* = 0) and moving in the direction it is oriented (*ψ* = 0). For simplicity, the following analysis uses *k_ϕ_* = *k_ψ_* = 1; the same qualitative results hold for different choices of *k_ϕ,ψ_* (see Supplemental Materials and Fig. S1). To understand how these control laws would affect the fly’s movement relative to the ambient wind direction we introduce a new variable for the course direction relative to the ambient wind direction,

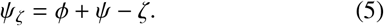

Next we calculate the derivative of *ψ_ζ_*, under the assumption that the fly changes its course direction much faster than any changes in the ambient wind direction (i.e. 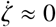),

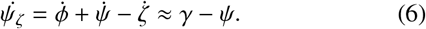

We now rearrange Eqn. 5 as *ϕ* – *ζ* = *ψ_ζ_* – *ψ* and substitute it into Eqn. 3, followed by a substitution of 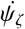 for the two instances of *γ* – *ψ* yielding the following,

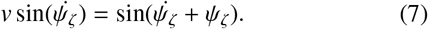

We now solve Eqn. 7 for 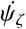 as follows,

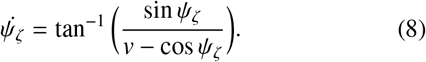

This expression allows us to analyze the stable and unstable equilibria for global course direction in relation to the ambient wind direction for different values of *v* using a 1-dimensional phase portrait (Fig. 2). This analysis shows that although moving downwind and upwind are both equilibria, upwind movement (*ψ_ζ_* = ±*π*) is the only stable outcome.

**Fig. 2:**
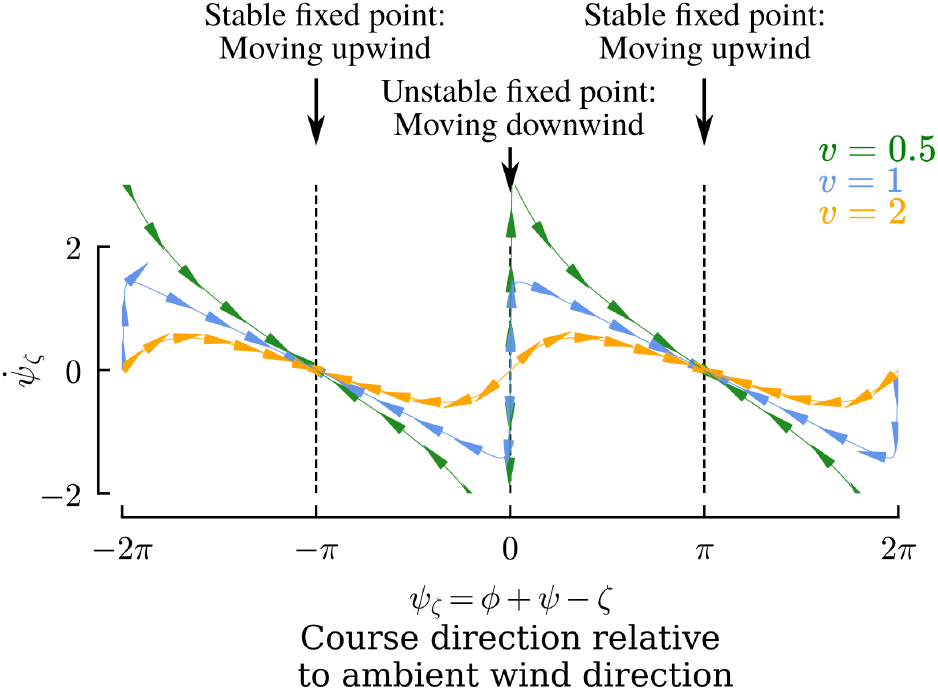
Visual anemotaxis guarantees stable upwind movement regardless of *v*. Each trace shows the phase portrait for different values of *v* given Eqn. 8. The arrows provide a visual aid that shows the flow along the phase space; their direction is determined by the sign of 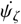.

During behaviors such as migration, insects may choose to fly at an angle relative to the wind [25], rather than orient upwind. Such behavior could be accomplished using visual anemotaxis, but the insect would not have access to what this angle is by using visual anemotaxis alone (see Supplemental Materials and Fig. S2).

### B. Active Anemosensing

In contrast to visual anemotaxis we now develop a framework for estimating the ambient wind direction that does not first require finding upwind. To begin, we simplify the measurement space by introducing two new variables (*α*: direction of air velocity relative to the global frame, and *δ*: “slip” angle between the air velocity and ground velocity vectors), and rewrite Eqn. 3,

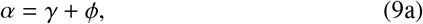

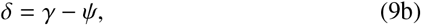

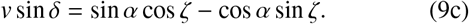

Eqn. 9 has two unknowns *v* and *ζ*, resulting in an under-determined system given the measurements *α* and *δ*. To uniquely determine *ζ*, we add the following constraint from the time-derivative of Eqn. 9,

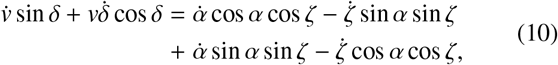

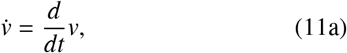

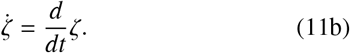

Together, equations 9c-11 fully constrain *ζ* provided that derivatives of the measurements can be calculated or mea-sured. Thus, in principle a variety of optimization methods could be used to estimate *ζ* using these relationships as constraints. To understand which trajectories lead to accurate estimates under varying levels of sensory noise and for different natural wind dynamics we develop an optimal framework for estimating *ζ* using convex methods in the following section.

### C. Active Anemosensing using Convex Optimization

In this section we construct an objective function of the form ℓ = ∥*L*∥ + ∥*dL*∥ that is linear with respect to the unknowns such that it can be efficiently solved using traditional convex optimization tools.

The terms in Eqn. 10 containing 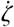 include the product of two unknowns, resulting in a non-convex formulation. Thus, we introduce the assumption the dynamics of the trajectory are faster than the dynamics of the wind. That is, for any given time window of length *τ* we assume that 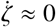, but the other derivative terms may all be non-zero and time-varying.

We can now build a convex estimator that incorporates measurements from a given window of time [*k* – *τ, k*] to estimate the average wind direction for that window 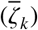. To build our estimator we define the following two loss functions derived from Eqns. 3,10:

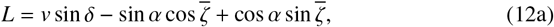

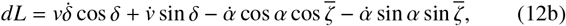

which define a fully constrained convex optimization problem over the time window [*k* – *τ, k*] to estimate the average ambient wind direction 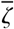:

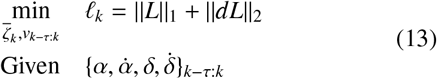

This optimization problem will find the best estimate for a single value of 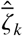, and a timeseries for 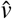, given sensory measurements from a window in time from *k–τ* to *k*. Because the errors in the derivative estimates will almost always be larger than the errors in the undifferentiated values, we apply a 2 – *norm* to *dL* and a 1 – *norm* to *L*. Although the optimization problem given in Eqn. 13 is convex, it does not follow Disciplined Convex Programming (DCP) rules because of the trigonometric functions. To efficiently solve the problem we can recast it as two DCP problems. For the first, we divide *L, dL* by cos 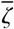, absorbing this constant term into *v*, denoted as *v^c^*, and then optimizing to find tan 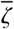 using the following expressions:

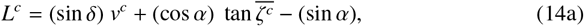

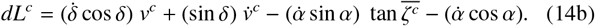

Extracting 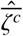 from tan 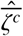 results in an ambiguity of ±*nπ*, 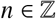, which we resolve using the following constraint derived from Eqn. 9:

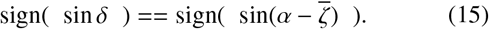

If 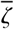 happens to be close to *π*/2 ± *nπ*, 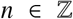, then dividing by 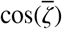 can result in large errors. Thus, to improve accuracy we solve a second optimization problem where we divide by 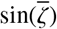 instead of 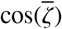 and optimize to find 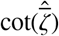 using the following modified expressions:

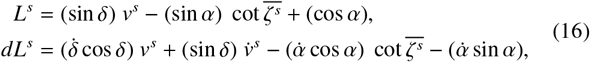

and choose the value of 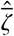 from the choices 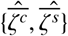 that corresponds to the smaller of the two values for 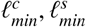. This results in choosing the value of 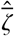 that is most consistent with the sensor measurements over the given window. Note that because we estimate an average wind direction given measurements from a segment of time *τ*-long, the estimate 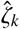 is delayed by *τ*/2. In our analysis we automatically adjust for this delay in order to compare the estimate to the true values, however, in a real-time application smaller values of *τ* should be preferred.

### D. Practical Considerations and Time Constants

To mitigate the effects of noise we assimilate measurement data across time using three time constants (Fig. 3A). These time constants are fundamental to our overall active anemosensing hypothesis, and not a specific feature of the convex solver we implement. With respect to our convex solver, the time constants define the window over which the solver is defined to operate. Therefore, the impact of these time constants on the resulting wind direction estimates should generalize beyond our particular convex solver.

**Fig. 3:**
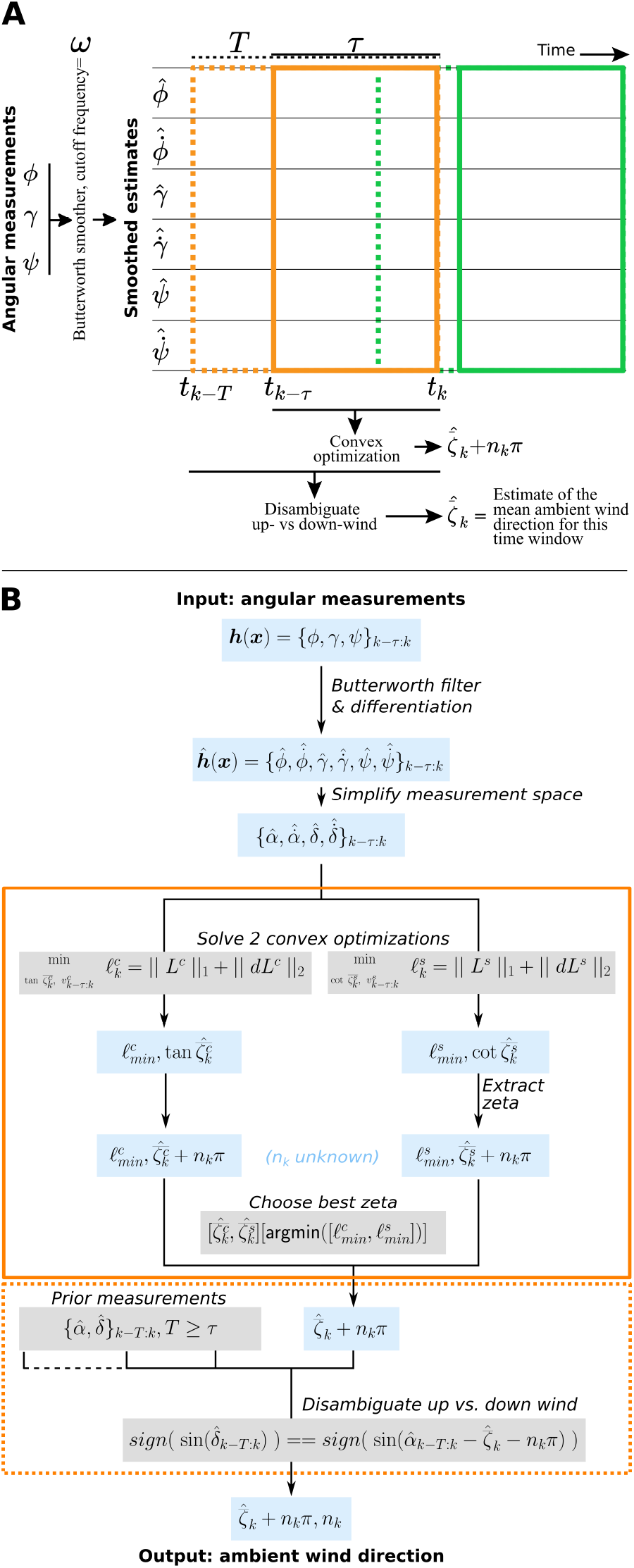
(A) Visual representation of the three time constants (*ω, τ, T*) fundamental to our active anemosensing hypothesis. (B) Outline of our convex ambient wind direction estimation algorithm. Note that the convex optimization steps in the orange box could be replaced with other workflows that are less optimal but more computationally efficient by using simplifications and heuristics, as we discuss in the Discussion.

(1: *ω*) We estimate smoothed values for the measurements 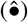, and their derivatives 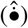, using a butterworth filter with a specified cutoff frequency, *ω* [26] to reduce noise in the estimates. The cutoff frequency effectively describes the maximum frequency of expected changes in the measurement. Since our algorithm assumes that changes in direction of motion occur at a higher rate than changes in wind direction, a good choice is generally an *ω* = turning frequency.

(2: *τ*) To estimate the ambient wind direction, we assimilate sensory measurements over a window in time of size *τ*.

(3: *T*) We find that resolving the *nπ* ambiguity (Eqn. 15) sometimes benefits from incorporating measurements across a longer time span, *T*, compared to the time span *τ* used for estimating *ζ* ± *nπ*.

Figure 3B provides a visual summary of our entire convex estimation framework, taking these three practical considerations and time constants into account.

## III. Methods

This section provides an overview of our implementation of the convex estimation framework described in the prior section, real world wind measurements, candidate trajectory selections, simulations, and our drone platform for validating the robustness of our estimator framework.

### A. Implementation of Estimation Framework

To solve the convex optimization problems outlined in Fig. 3 we used the open source python package pynumdiff to estimate butterworth smoothed derivatives [27], and cvxpy [28] with the proprietary MOSEK solver [29]. For computational efficiency we estimated wind directions for every 10^th^ time step (6 Hz for our validation experiment and 1 Hz for our simulation experiments).

### B. Real world wind measurements

We collected wind direction and speed data using two 3-D ultrasonic anemometers (Trisonica mini, Anemoment, Longmont CO) operating at 10 Hz positioned orthogonal to one another to ensure accurate readings of both horizontal and vertical wind components. We collected data in an open desert environment (Fig. 4A). The vertical wind component was minimal, whereas the horizontal wind varied, with dynamic periods characterized by low speeds and frequent changes in direction, as well as consistent periods with higher wind speeds and near-constant direction (Fig. 4B).

**Fig. 4:**
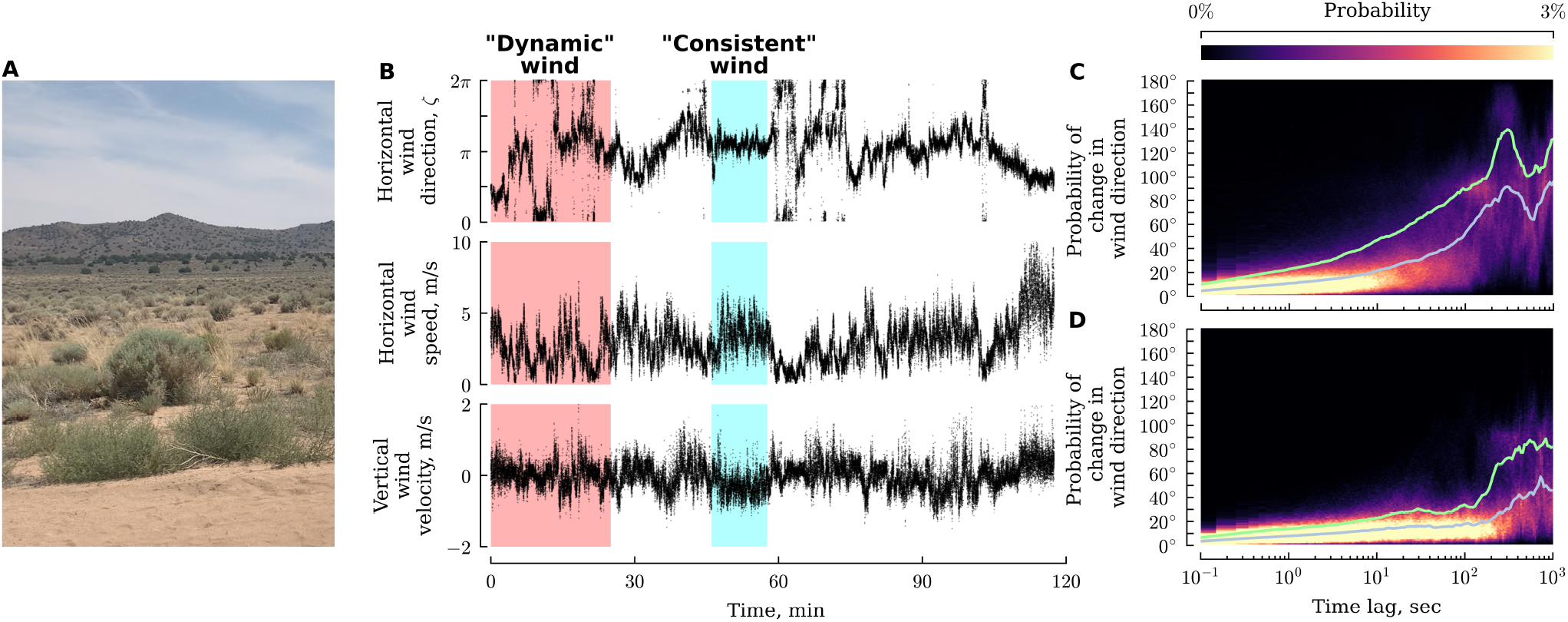
Natural wind exhibits periods of both dynamic or consistent wind directions. (A) Landscape where wind measurements were collected in northern Nevada. (B) Time series of wind direction and speed. Shaded regions indicate segments used throughout our analysis. (C-D) Probability density of changes in wind direction as a function of elapsed time for the two shaded regions shown in B (see text for detail).

To visualize the likelihood of a shift in wind direction over time, we took the difference between a point in the recorded time-series and a point at various time lags between 0.1 and 1000 seconds. The differences in wind direction at each lag were computed by moving both forward and backward in time, and those values were averaged. By repeating this analysis throughout the time-series, we created a frequency distribution. We then used this distribution to generate a heat map, which demonstrates the approximate probability and magnitude that wind direction will change at any given time over a period of 1000 seconds. In our dynamic wind segment, changes in wind direction became increasingly likely after 10-50 seconds (Fig. 4C), whereas for the consistent wind segment large changes in wind direction were rare for the entire time series (Fig. 4D).

### C. Candidate trajectories

To design our candidate trajectories, we took inspiration from previously published data on plume tracking fruit flies flying in a laminar flow wind tunnel [22]. In the presence of attractive odors, fruit flies perform upwind and crosswind flight maneuvers referred to as surges and casts (Fig. 5A).

**Fig. 5:**
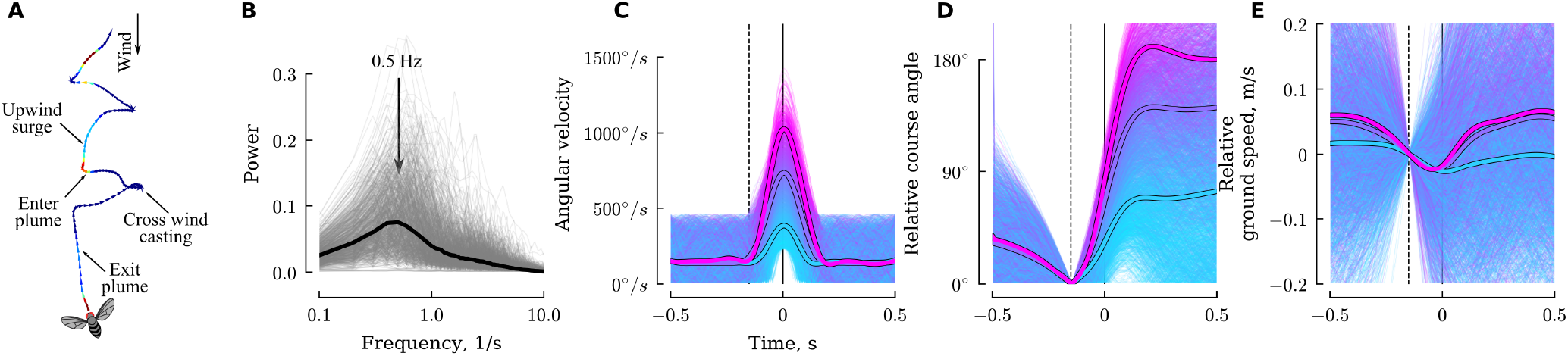
Plume tracking flies initiate turns at 0.5 Hz and decelerate as they enter the turn. (A) Stereotypical flight trajectory of a plume tracking insect from [22]. (B) Power spectra of of the cross-wind velocity for trajectories during which flies experienced an attractive odor, are moving on average upwind, and are at least 1 second long (N=700 trajectories). (C) Angular velocity during isolated turns from the trajectories in B (n=4006 turns). Color encodes peak angular velocity. Thick lines show the average for three groups corresponding to peak angular velocity ranges of [230° – 570°], [570° – 917°] and [> 917°]. (D) Change in course angle relative to the point in time indicated by the dashed line. (E) Change in ground speed relative to the point in time indicated by the dashed line.

Although this behavior has received substantial attention, aspects of their flight dynamics have escaped scrutiny, which we briefly address here by reanalyzing these previously published data. To focus our analysis on plume tracking individuals, we selected trajectories with durations greater than 1 second during which an odor (ethanol) was encountered at least once while moving (on average) upwind in 0.4 m/s wind, resulting in 700 trajectories. These flies exhibited regular changes in course direction at a rate of approximately 0.5 Hz (Fig. 5B), corresponding to ~ 1 left and ~ 1 right turn every two seconds.

Flies generally accomplish changes in course direction using rapid turns called saccades [19]–[21], [30]. To understand the dynamics of the saccades during plume tracking specifically, we isolated individual turns from the 700 trajectories by selecting 1-second long snippets during which the flies exhibited a minimum angular velocity of 230°/sec, and no greater than 460°/sec prior or after the turn (Fig. 5C). This threshold choice is consistent with prior saccade analyses of freely flying flies which used a threshold of 300°/sec [19]. The amplitude of their turns range from small adjustments to 180° changes (Fig. 5D), with an overall average of 110°. As with flies flying in the absence of odor or wind [19], the saccade magnitude is strongly correlated with the peak angular velocity during the turn, suggesting they have an internal model of their new desired heading prior to initiating the turn, an observation that goes against the prediction of the visual anemotaxis hypothesis. In order to make these rapid changes in direction flies first decelerate, and then accelerate as they exit the turn (Fig. 5E), though there is substantial additional variability in velocity throughout their trajectories.

Based on the behavior of plume tracking flies, we primarily consider trajectories stereotyped by regular changes in direction accompanied by synchronized changes in ground speed. We consider three categories of trajectories with small (~20°), medium (~90°), and large (~160°) turn angles. For the small turn trajectories, the magnitude of each turn is selected randomly from a uniform distribution of 16-24°. For medium and large turn trajectories these uniform distributions were 72-108° and 144-171°, respectively. We applied a Gaussian smoothing kernel to these piecewise constant course directions, resulting in smooth yet still concentrated turns. Although plume tracking insects do exhibit vertical movements, we restricted our candidate trajectories to movements parallel to the ground plane for simplicity. For the orientation of our simulated flies, we consider two cases inspired by the active control required by visual anemotaxis: either constant-*ψ* 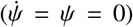 or constant-*γ* 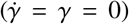. Finally, we also consider constant ground speed trajectories.

### D. Simulations

To determine the accuracy of ambient wind estimates for different trajectories in natural wind conditions, we simulated our trajectories in both the 25 minute long dynamic- and 12 minute long consistent-wind scenarios at a temporal resolution of 0.1 seconds (the time resolution of our wind data). To focus our analysis on the impact of the trajectory shape, rather than the feedback control necessary to achieve that trajectory, we programmed the trajectories kinematically, thus changes in wind speed and direction did not have any effect on the shape of the trajectories in our simulations. We justify this decision by noting that flies exhibit exceptional visual and mechanosensory feedback mechanisms for gust rejection [31]. We extracted the three sensory modalities under consideration, and applied zero-mean Gaussian noise with standard deviations ranging between 17° – 67°. All simulations were performed with the same initial random seed to ensure a fair comparison across different conditions.

Next we smoothed the sensor measurements and estimated their derivatives using a butterworth filter and ran the estimates through our optimal estimation framework for different values of *τ* and *T*. For each individual wind direction estimate, we calculate the absolute value of the error between the estimate and the true wind direction.

### E. Drone hardware for validation experiments

#### Platform

To validate our estimation approach we built a quadrotor drone using a Holybro X500 frame (Fig. 6A). The drone was controlled by a PixHawk flight controller. The PixHawk was flashed with PX4 software and configured using QGroundControl software. The drone was powered by a 14.8 V 5200 mAh 60C 4S LiPo battery (Zeee Power), which provided approximately 5-10 minutes of flight time with our full payload in ~ 2 m/s wind. The drone was piloted manually, using a 2.4GHz controller (FrSky Taranis Q X7) that communicated flight commands to a compatible receiver (FrSky Taranis Receiver X8R) connected to the PixHawk.

**Fig. 6:**
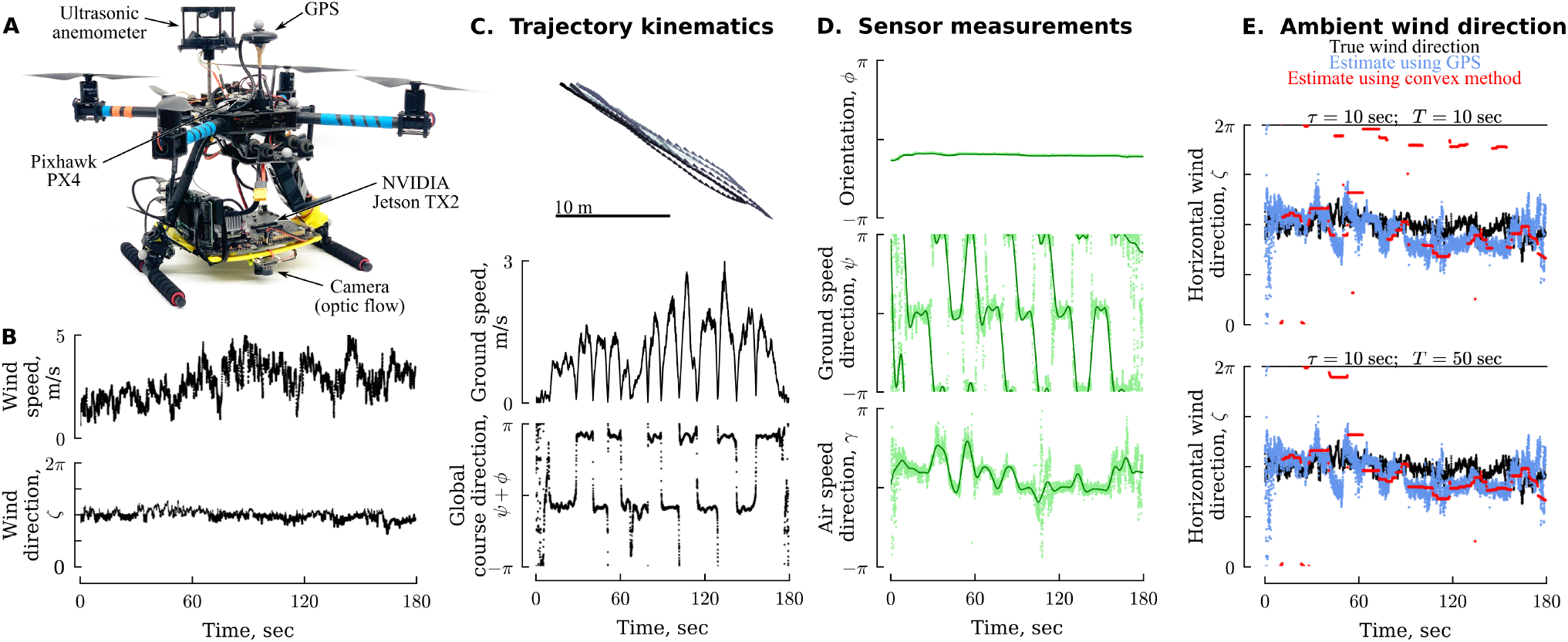
Ambient wind direction estimation using the framework outlined in Fig. 3 is robust to real world sensor data. (A) Drone used for data collection. (B) Ambient wind direction and speed during piloted flight. (C) Trajectory kinematics of piloted flight provided by GPS measurements. (D) Sensor measurements (light green points) and smoothed sensor estimates (dark green lines). (E) Comparison of true ambient wind direction repeated from B (black), estimated wind direction using GPS derived ground speed measurements (blue), and estimated wind direction using the sensor data from D and the framework outlined in Fig. 3 for two different values of *T* (red). Convex estimates are aligned based on the median time stamp in a given window for *τ*.

#### Sensors

The PixHawk board has a built in IMU, which provides heading and attitude angles, angular velocities, and linear acceleration measurements at 60 Hz. To provide ground truth data for our experiment we mounted a 60 Hz GPS (Holybro M8N GPS) to the drone. Validating our estimation protocol required additional sensors for optic flow (providing *ψ*) and air speed (providing *γ*). For optic flow we mounted a USB camera (ELP 1080p camera with a 2.9mm wide angle lens) underneath the drone operating at 30 Hz. Image data was used to calculate 2D optic flow in real time using the Lucas Kanade algorithm [32], [33] with OpenCV [34]. For air speed measurements we mounted a 3D ultrasonic anemometer (Trisonica mini from Anemoment) to the top of the drone operating at 40 Hz.

#### Data collection

To collect sensor data, we installed an NVIDIA Jetson TX2 equipped with a a 1 TB NVMe M.2 SSD below the drone. We flashed the TX2 with Linux4Tegra and installed ROS (Robot Operating System) to provide a centralized interface to all of our sensors. We used the MavROS package to interface with the PixHawk, which provides access to the IMU measurements and RC controller inputs. Two other custom ROS packages were used to calculate optic flow and interface with the anemometer. All flight data was recorded into a single ROS bag file for offline analysis. All sensor data was linearly interpolated to the highest sensor frequency of 60 Hz prior to analysis, along with corrections for declination and sensor orientations.

## IV. Validation

To verify our algorithm’s robustness to real world data we manually piloted a sensor-equipped drone in a designated radio controlled flight space (Reno Radio Control Club Field) in the northern Nevada desert (Fig. 4A). We recorded the ambient wind direction and speed with a pair of orthogonal ultrasonic anemometers (Trisonica Mini, Anemoment, Longmont CO) positioned approximately 2 m off the ground. We manually piloted the drone back and forth at ~ 1 – 3 m/s (Fig. 6C) at an altitude of ~ 4 m, and recorded the three sensor modalities of interest (Fig. 6D). For our butterworth smoothing filter we chose a cutoff frequency of *ω* = 0.1 Hz (the approximate turning frequency of the drone’s trajectory), which helped to reduce the inherent sensor noise.

To provide a ground truth for the ambient wind direction estimates, we vector-subtracted the GPS provided absolute ground velocity from the two horizontal components of the apparent airspeed (Fig. 6E, blue). This estimate provides the best-case scenario for any 2-dimensional wind direction estimator that relies on the airspeed measurements, allowing us to decouple the performance of the estimation algorithm and the choice of sensors. These GPS and airspeed derived estimates do deviate from the ground truth (median error of 21°). Some of these errors are explained by vertical wind measured by the drone-mounted anemometer (linear regression *R*^2^ = 0.1, *pval* < 10^-3^). Since the ambient wind exhibited very little movement in the vertical dimension, and the drone exhibited very little pitch and yaw during its trajectory, the changes in vertical wind measured on the drone were likely due to changes in downdraft from the rotors required for stabilizing the altitude of the drone. Smaller flying systems that introduce less downdraft to maintain lift (such as an insect) may be less impacted by this effect. Alternatively, future efforts will be needed to ensure a more accurate sensor estimates of airspeed direction.

Finally, we applied our optimal estimation algorithm to the sensor data to estimate the ambient wind direction with *τ* = 10 sec (the approximate turning frequency of the drone), and two values of *T* (Fig. 6E, red). The larger value of *T* = 50 provided better discrimination between the up- and downwind options. Our optimal estimation approach using only angular sensor measurements compares favorably with the ground truth estimate using the GPS and airspeed measurements (median error of 9.5° for *T*=50 sec case). These results show that the algorithm outlined in Sec. II that relies purely on angular direction information is sufficiently robust to handle real-world data to provide wind direction estimates on par with other methods that also take advantage of magnitude information.

## V. Results

Using a series of simulations we set out to determine which types of trajectories can provide the most accurate ambient wind direction estimates. For each candidate trajectory we consider (e.g. Fig. 7A-B), and for a variety of sensor noise magnitudes, we calculated the corresponding sensor measurements for either the dynamic or consistent wind scenario, and estimated the smoothed sensor estimates and their derivatives for a given choice of *ω* (e.g. using a cutoff frequency *ω*=turning frequency: Fig. 7C; a detailed analysis of *ω* is given in the following section). Then, for a given choice of *τ* and *T* we estimated ambient wind direction values using our convex optimization framework (e.g. Fig. 7D) and calculated the associated errors (e.g. Fig. 7E). These errors largely stem from rapid changes in wind speed or direction, insufficient changes in course direction per unit *τ*, and sensor noise, as we will show in detail in subsequent analyses. Repeating this analysis for a variety turning frequencies and values of *τ* revealed that there is an optimal choice of *τ* for any given turning frequency (Fig. 7F). In general, we find that choosing *T* = *τ* works well, however in some cases, particularly for small values of *τ*, choosing *T* > *τ* provides a modest reduction in the error compared to *T* = *τ* (Fig. 7F, stars), suggesting that distinguishing between up- and down-wind is a particular challenge (the following section will elaborate on the choice of T). This is also apparent in the bimodal distribution of errors seen in some scenarios (e.g. turning frequency=0.02 Hz, *τ* = 10, 50 sec).

**Fig. 7:**
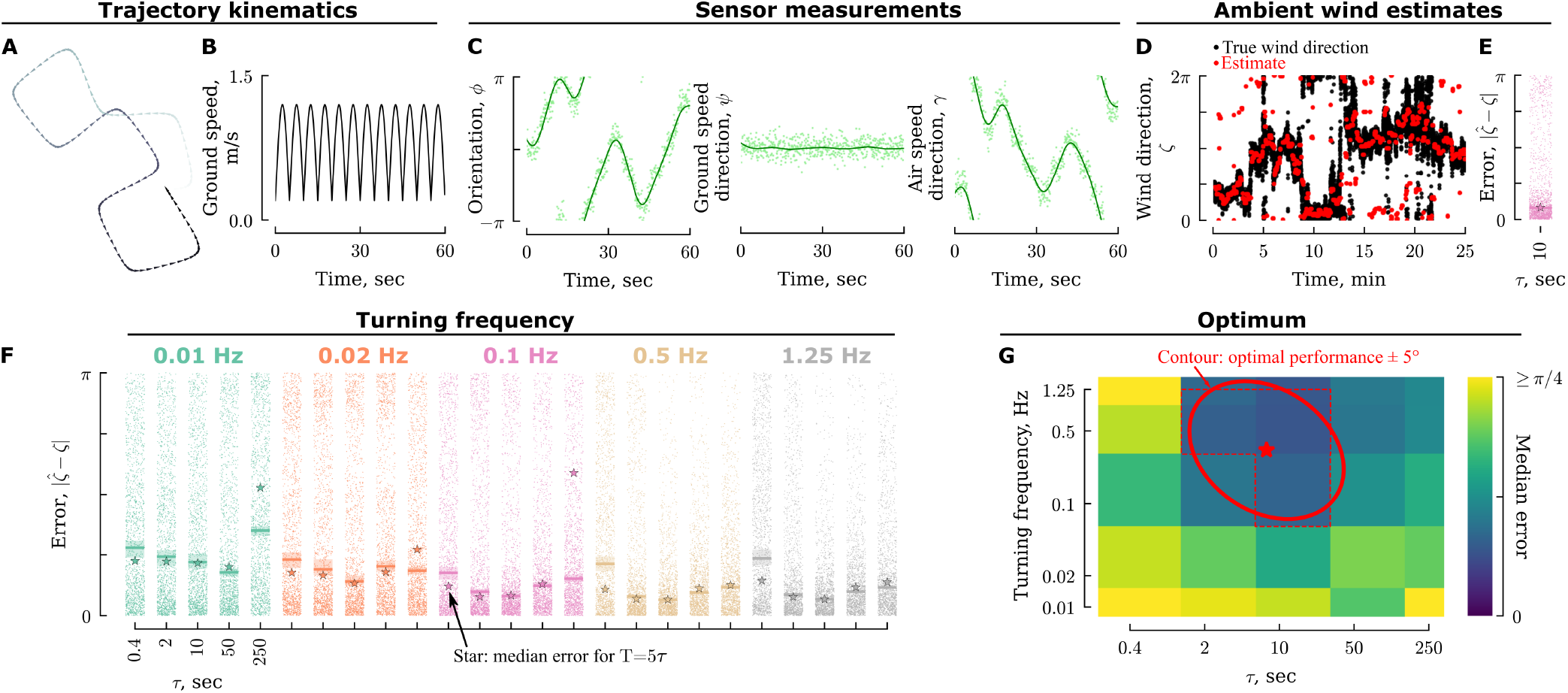
(A) Example simulated trajectory and (B) ground speed for a turning frequency of 0.1 Hz. (C) Sensor measurements for the trajectory shown in A-B simulated in our dynamic wind scenario (Fig. 4B, red shading) with 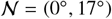 Gaussian noise applied (light green dots), along with subsequently smoothed estimates (dark green lines). (D) Comparison of the true ambient wind direction and estimate for *τ* = *T* = 10. (E) Error in wind direction estimate for each estimate shown in D, along with the median and 95% confidence interval. Star: median error for *T* = 50. (F) Simulation results for different combinations of turning frequency, *τ* and *T*, plotted as in E. (G) Median error values from F plotted as a heatmap with respect to turning frequency and *τ*. Red dashed lined: contour indicating the best performance ±5°. Ellipse: best fit ellipse representing the red dashed contour. Star: center of ellipse.

To better visualize the optimal choice of both *τ* and the turning frequency, we plot the median errors for each case as a heatmap (e.g. Fig. 7G). In many of the cases we consider, several combinations of *τ* and the turning frequency offer similar performance. To capture this range of optima, we determined which combinations result in a median error that is within ±5° of the optimum (Fig. 7G red contour), and approximate this contour with an ellipse for visual simplicity. From this analysis we can conclude that for trajectories with 90° turns, variable velocity, and constant *ψ* = 0, in our dynamic wind scenario with sensors subject to modest noise, the optimal turning frequency ranges from 0.1-1 Hz, with values of *τ* ranging from 2-50 seconds.

### A. Analysis of time constants

The optimal combinations of turning frequency and *τ* all occur where there is at least 1 turn per time *τ* (Fig. 8A). At sufficiently long values of *τ*, however, there is a high likelihood of the wind having changed direction in our dynamic wind scenario, which increases the median error. As a result, the optimal choice of *τ* is constrained to be large enough to capture a change in flight direction, but short enough to avoid likely changes in ambient wind direction. The changes in flight direction, meanwhile, must occur more frequently than changes in wind direction. In our dynamic wind scenario the wind begins to show a high likelihood of changing direction after 10-50 seconds (Fig. 8A, blue dashed line), thus turning frequencies of 0.1 Hz or greater are required to achieve small errors.

**Fig. 8:**
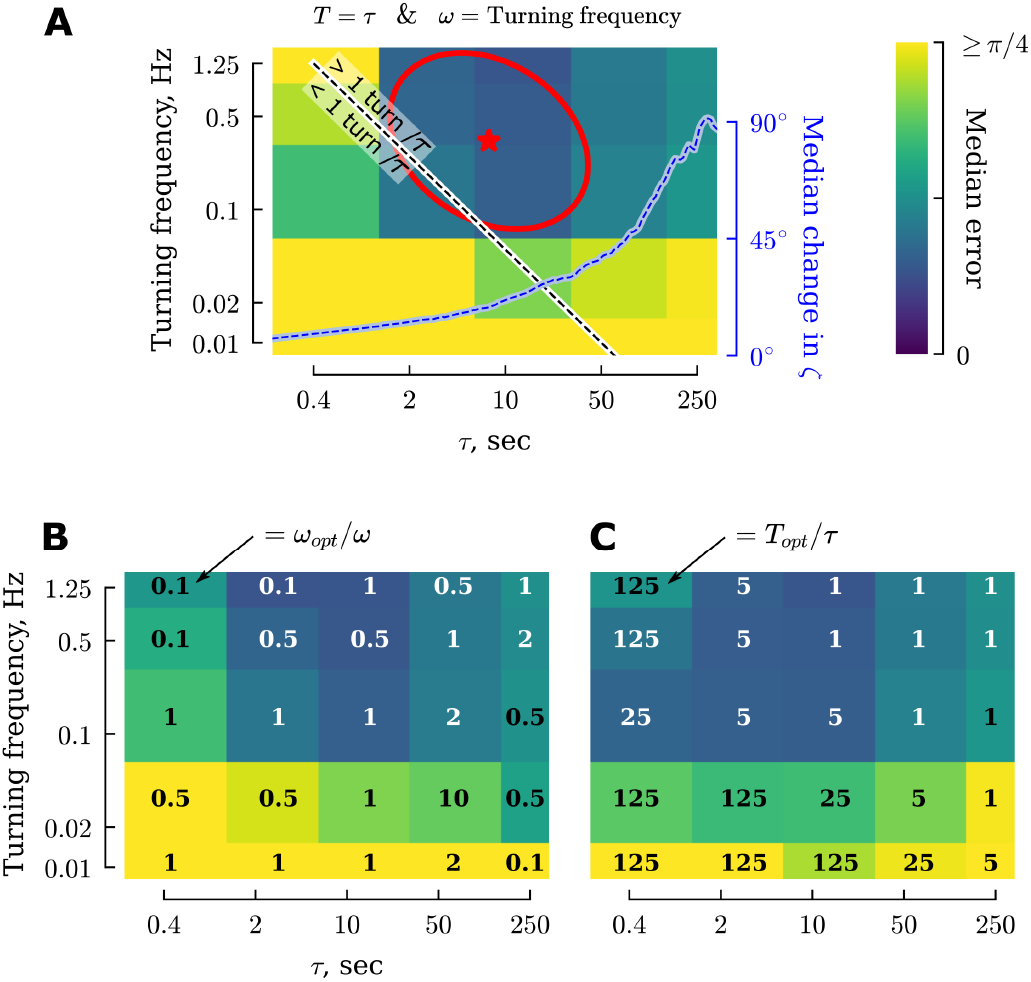
Turning frequency dictates optimal choices of *ω, τ*, and *T*. (A) Heatmap duplicated from Fig. 7G, i.e. dynamic wind, 90° turns, and Gaussian sensor noise 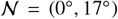. Combinations of turning frequency and *τ* above the white-black line correspond to cases where there is at least on full left or right turn per unit *τ*. The blue dashed line shows the median probability of a change in wind direction for our dynamic wind scenario, copied from Fig. 4C. (B) Heatmap plotted as in A, but showing the lowest median error across five choices of *ω*; numbers indicate the optimal choice of *ω_opt_* relative to a nominal choice of *ω*=turning frequency. (C) Heatmap plotted as in A, but showing the lowest median error across four choices of *T*; numbers indicate the optimal choice of *T_opt_* relative to *τ*. For good performance, *T* is always sufficiently large to encompass at least one left or right turn.

To better understand the sensitivity of our results to choices of the cutoff frequency *ω* in our smoothing filter, we performed a parameter sweep of *ω* (Fig. 8B). We considered values of *ω* equal to the product of the turning frequency and five constants: [0.1, 0.5, 1, 2, 10]. For combinations of fast turning frequencies and short *τ* (upper left corner of the heatmap), choosing a smaller cutoff frequency (0.1*turning frequency) provided a modest reduction in median error of the wind direction estimates. Overall, however, a choice of *ω* =turning frequency generally performs well. This makes sense as the majority of changes in the sensory measurements are due to changes in course direction of the trajectory.

Next we performed a parameter sweep of *T*. We considered values of *T* equal to the product of *τ* and four constants: [1, 5, 25, 125]. For combinations of fast turning frequencies and short *τ* choosing *T* = *τ* performs poorly (Fig. 8A). However, choosing a much larger value of *T*, e.g. *T* = 125*τ*, substantially improves the accuracy of ambient wind direction estimates for these scenarios (Fig. 8C). For a turning frequency of 0.5 Hz and *τ* = 0.4 sec the optimal choice of *T* = 125*τ* = 50 sec, which incorporates information across 50 individual turns. Overall, we conclude that assimilating sensory measurements across at least one turn is required, but is most important for distinguishing between up and downwind.

### B. Analysis of turn angle & sensor noise

Next we set out to answer the question: are small or large turns more effective for estimating the ambient wind direction under dynamic wind conditions? We repeated the analysis summarized in Fig. 8A for trajectories consisting of small, medium, and large turns (Fig. 9A). Across a wide range of sensor noise, larger turns consistently provide more accurate ambient wind direction estimates. As with the first example provided, across all cases the lowest errors are achieved when there is at least one turn per unit *τ*. Choosing *ω*=turning frequency again works well across this entire range of turn angles and sensor noise levels (Fig. S3). For short *τ*, estimation errors can again be reduced by choosing large enough values of *T* to incorporate measurements across at least one turn (Fig. S4).

**Fig. 9:**
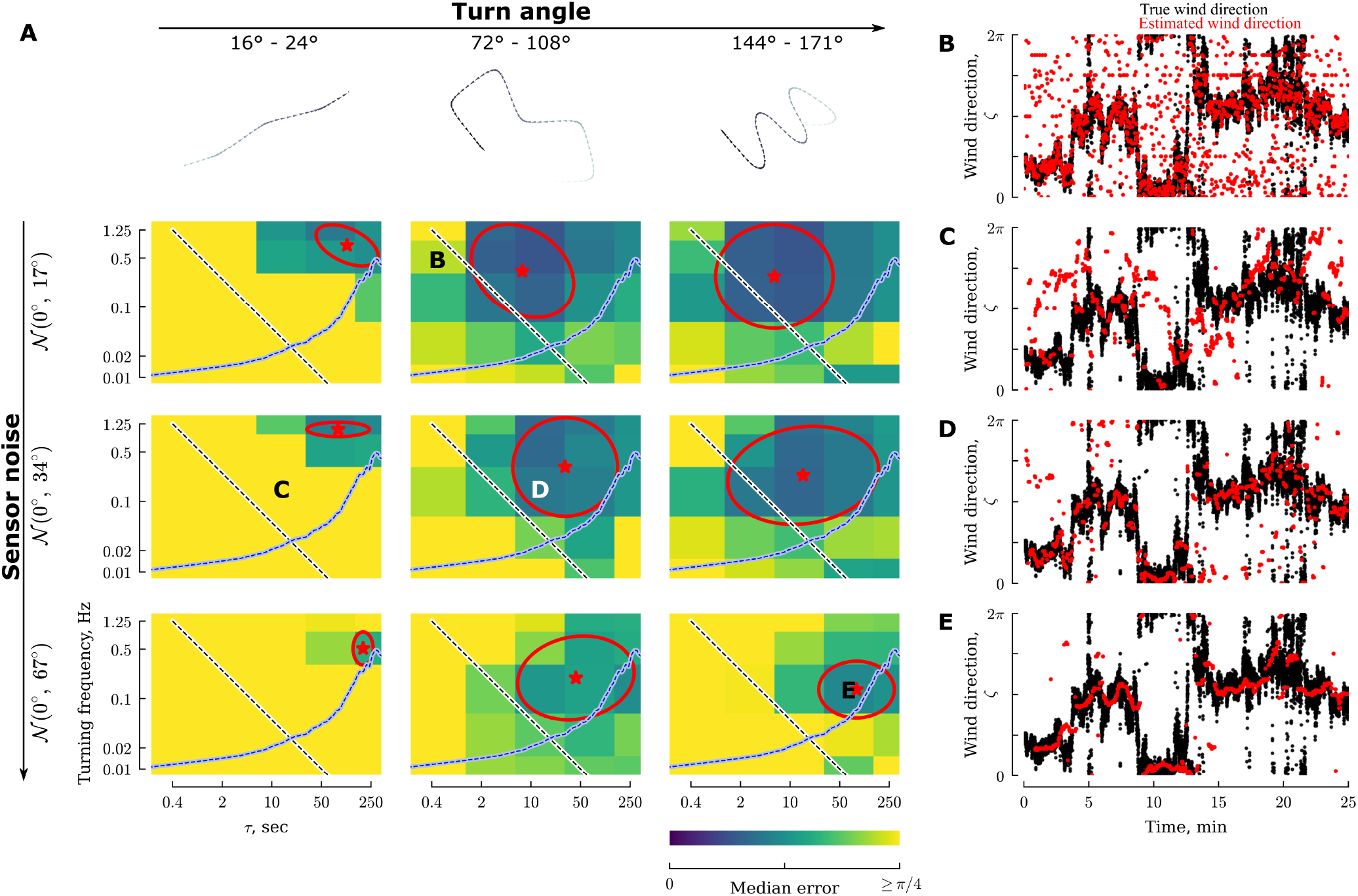
Larger turn angles result in better performance and smaller optimal values for *τ*. (A) Simulation results for the dynamic wind segment plotted as in Fig. 7G. (B-E) Representative examples comparing true (black) and estimated (red) wind directions for combinations of turn angle, noise, turning frequency, and *τ* as indicated by the letter placements in A.

To provide some intuition about how accurate or poor the estimates of ambient wind direction can be, and characteristics of the errors, we show four representative examples in Fig. 9B-E. For frequent turns and short *τ*, the estimates are noisy, but the largest errors are characterized by errors in distinguishing between up- and down-wind, i.e. errors of ±*π* (Fig. 9B). Small turn angles result in poor estimates even when information is assimilated across at least one turn (Fig. 9C). For large turns, assimilating measurement information across a larger value of *τ* results in smoother (less responsive) estimates (compare Fig. 9D-E).

For larger values of sensor noise, regardless of the turn angle magnitude, the optimal range of turning frequency decreases slightly. One likely explanation for this is that as the sensor noise increases it becomes increasingly difficult to filter out noise without sacrificing the signal. Slower turning frequencies help to alleviate this by allowing the use of a more aggressive cutoff frequency that can smooth the noise more effectively. To maintain at least one turn per unit *τ* for these slower turning frequencies, however, means that the optimal value for *τ* must increase. This makes the estimator less able to respond to rapid fluctuations in wind direction (e.g. Fig. 9E).

There are essentially four categories of errors lumped together under our single error metric: (1) poor estimates of 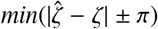 (i.e. the errors that are (much) less than *π*), which typically result from either too small a choice of *τ* or too small a turn angle (e.g. Fig. 9B-C);(2) poor estimates of the ±*nπ* ambiguity, typically resulting from too small a choice of *T* (compare Fig. 8A,C); and (3) overly smooth estimates of 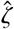 as a result of integrating sensory information over too large of a time window *τ* (e.g. Fig. 9E); (4) poor estimates of 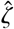 resulting from rapid changes in the ambient wind direction over the course of *τ*. To mitigate all error categories requires an intermediate value of *τ*, medium to large turn angles, and a sufficiently large value of *T*.

### C. Dynamic vs. consistent wind

Next we turn our attention to how the optimal values of *τ*, turning frequency, and turn angle change if the wind has a more consistent direction over time (Fig. S5). In contrast to the dynamic wind case (where larger turn angles help to reduce the median error in 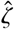), we find that in consistent wind the overall error achieved by trajectories with small, medium, and large turns is similar (Fig. 10), suggesting that large turns are particularly important when the wind direction is dynamic. Meanwhile, consistent wind is less sensitive to the specific choice of turning frequency or *τ*. That is, larger values of *τ* as well as slower turning frequencies are both effective.

**Fig. 10:**
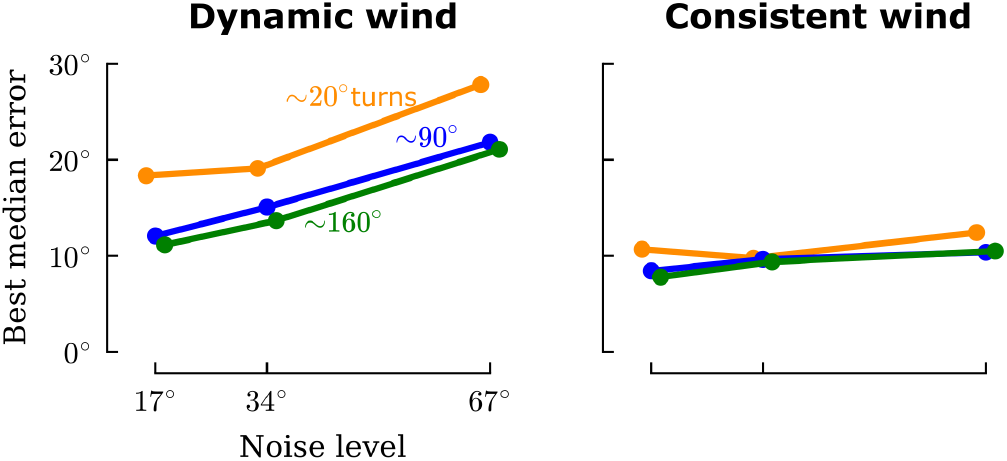
Median error in 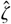 for trajectories with different turn angles in both the dynamic and consistent wind scenarios, as a function of zero-mean gaussian sensor noise with varying standard deviations.

To help summarize and generalize our results across a broader range of wind scenarios (Fig. 11), we individually analyzed shorter segments from both the consistent and dynamic wind examples. We randomly selected 1000 segments of random lengths ranging from 1 to 1000 seconds. For each segment we characterized the wind by the maximum and mean change in speed and direction relative to the middle of the segment, providing a measure of how self-correlated the wind was over that period of time. We then determined the minimum 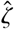 error for that particular segment, and the corresponding optimal *τ*’s and turning frequencies associated with that minimum error ±5° (i.e. the location and extent of the red ellipses shown in Fig. 9).

**Fig. 11:**
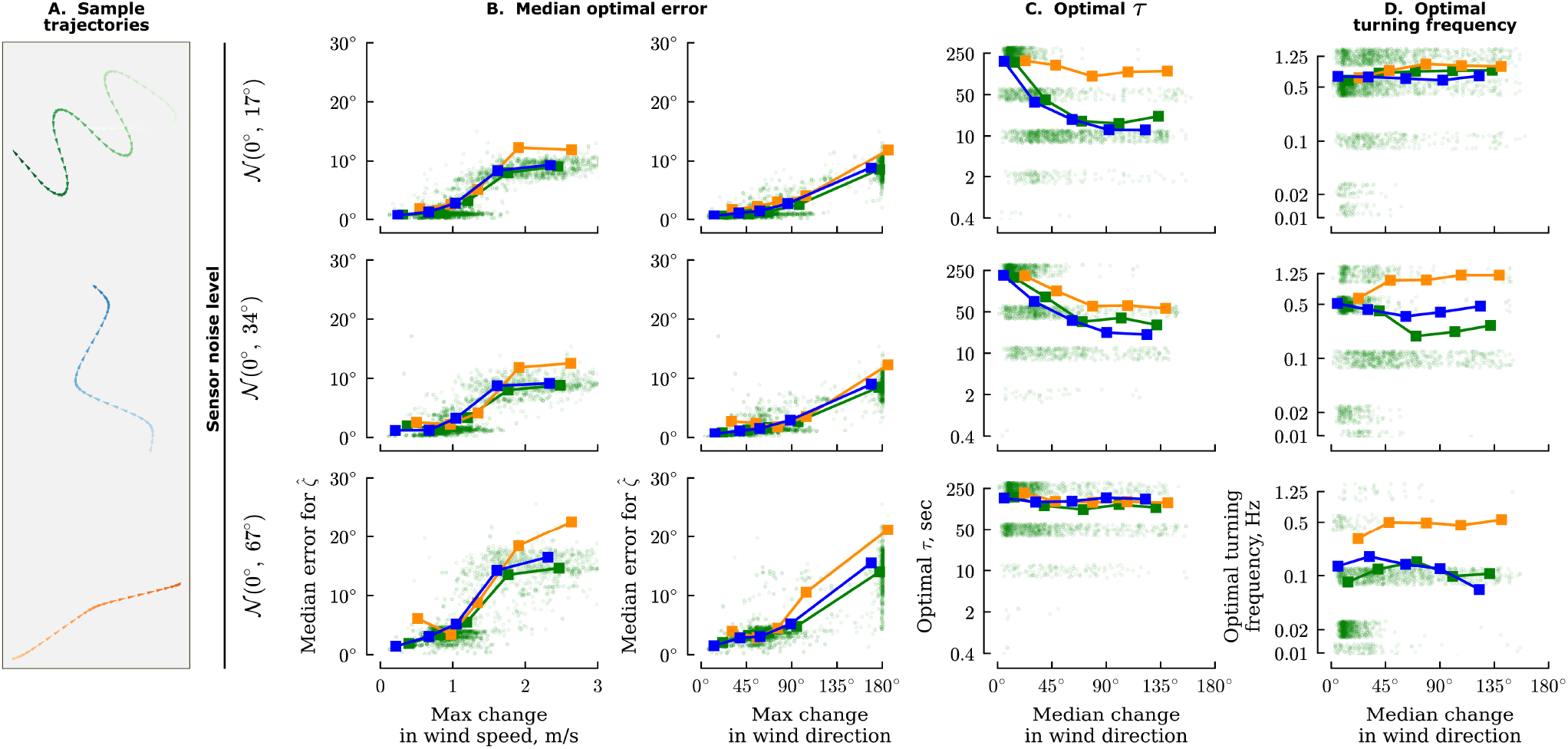
Summary of optimal error, *τ*, and turning frequency as a function of sensor noise, and wind variability for trajectories with different turn angles. (A) Example trajectories color-coded to the data in B-D. (B) Median optimal error for *ζ* as a function of the maximum change in wind speed and direction for 1000 different segments of wind ranging from 1-1000 seconds long. Light green points show the individual results for each segment for the large turn angle trajectories, bold squares show the median values for evenly spaced bins. (C) Optimal values for *τ* as a function of the median change in wind direction, plotted as in B. (D) Optimal turning frequencies as a function of the median change in wind direction, plotted as in B.

We find that errors in 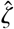 are strongly correlated with the maximum change in both wind speed and direction (Fig. 11B). The optimal choice of *τ* decreases for more dynamic wind segments allowing the algorithm to respond more quickly, but as the sensor noise increases, larger values of *τ* are better (Fig. 11C). Trajectories with small turn angles fare better with larger values of *τ*, which allows the estimator to incorporate information across a larger time window. This does, however, result in increased errors in the dynamic wind cases. The optimal choice of turning frequency remains relatively constant across different wind dynamics, though slower turning frequencies can also be optimal for very consistent wind (Fig. 11D). As sensor noise increases, the optimal turning frequency decreases, which makes sense since it becomes more challenging to distinguish between signal and noise.

### D. Constant ground speed & constant airspeed direction trajectories

Finally, we turn our attention to two variations of the trajectory candidates to answer two questions: would trajectories with constant ground speed 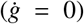 or constant airspeed direction (*γ* = 0) provide any improvements in the estimates over trajectories with variable ground speed and constant *ψ* = 0? These choices are motivated by the observation that many insects use visual information to maintain a constant ground speed [35], and typically orient their bodies in the direction of their airspeed [22]. The results for these trajectories are summarized in Figs. S6-S8/

Constant ground speed trajectories show almost no change in optimal median error, *τ*, and turning frequency compared to the variable ground speed trajectories. Constant *γ* = 0 trajectories, however, do show one subtle but important improvement: smaller values of *τ* under high levels of sensor noise (Fig. S6C). Why might this be the case? In our simulations the wind speed was typically higher than the ground speed (mean wind speed was 2.3 m/s for the dynamic wind scenario; mean ground speed was 0.84 m/s for a turning frequency of 0.1 Hz). Thus, by adopting a constant *γ* = 0 trajectory the agents and their sensors are generally oriented in a similar direction despite changing their direction of travel (compare the trajectory plots from Fig. S6). As a result, 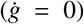 remains quite small for the constant *γ* trajectories, making it easier to accurately estimate *ϕ* with large levels of sensor noise (Fig. S9). With better estimates of the true sensor values, it is no surprise that the resulting ambient wind direction estimates are also more accurate.

## VI. Discussion

Recent release-and-recapture experiments with fruit flies in a feature-sparse desert showed that these tiny insects are able to locate the source of odor plumes from substantial distances over the course of ~ 10 minutes under a variety of wind conditions. In those experiments, flies were most successful at lower wind speeds [36]. Although this result is intuitive from a biomechanics perspective, these low wind speeds are typically associated with more variable wind directions [24], making the plume tracking task more challenging. Flies’ plume tracking success under such conditions suggests that they are adept at following odor plumes in variable direction wind conditions. Our active anemosensing hypothesis provides a potential explanation for how they might accomplish this by integrating three angular sensor measurements known to be encoded in their central complex: orientation, airspeed direction, and direction of movement.

### A. Summary of key results

We begin our discussion with a summary of our key results (see Figs. 11-S6). (1) Large changes in course direction are most effective for accurately estimating the ambient wind direction. This is especially true when the wind direction changes frequently. (2) For both dynamic and consistent wind cases, it is important that sensory information be integrated across at least 1 turn. (3) In more dynamic wind, shorter integration times (*τ*) are better, and faster turning frequencies are also necessary. (4) When subject to large amounts of sensor noise, slower turning frequencies offer better estimates by allowing the use of more aggressive smoothing filters to reject noise. (5) In terms of estimation accuracy, constant-ground-speed and constant-*γ* trajectories offer no substantial advantage over variable-ground-speed and constant-*ψ* trajectories. (6) For large amounts of sensor noise applied to trajectories with large turn angles, however, constant-*γ* trajectories allow for shorter integration times (smaller *τ*) compared to constant-*ψ* trajectories. (7) A key challenge is distinguishing between up- and downwind. In other words, it is challenging for an insect to determine if they are flying faster than the wind speed but headed downwind, versus flying slowly upwind. This challenge can be more reliably overcome by integrating information across a longer period of time (*T* > *τ*) to include information that spans more changes in course direction (see Fig. 8).

### B. Zig-zag maneuvers optimize wind direction estimation

The trajectories exhibited by flying and swimming animals during plume tracking across a wide range of taxa are nearly all characterized by some form of zigzagging, or casting [1]. Zigzagging, however, is not explicitly required for plume tracking if ambient wind direction information is directly accessible. For example, walking cockroaches are adept at following plumes but do not exhibit the classic zigzagging behavior [37]. Walking *Drosophila* also do not need to perform zigzagging trajectories when presented with a homogeneous and consistent odor plume, instead they will continue straight upwind [38]. For a walking animal, estimating the ambient wind direction from apparent wind estimates is trivial: they can either hold still for a brief moment, or rely on proprioceptive cues to accurately estimate their ground speed or distance travelled given their stepping speed and leg length [39], [40]. Indeed, visually blind flies have no trouble in finding the direction of ambient wind [38]. Of course, some changes in direction are required in order to stay centered on the plume, especially for turbulent plumes [41], but these changes in direction need not be rhythmic, frequent, nor large.

Furthermore, in some efforts to reverse engineer plume tracking strategies using an “optimal” reinforcement learning framework for agents that are provided with ambient wind direction information, the distinctive casting exhibited by animals appears to be largely absent [42], or diminished when the agent is inside the plume [43]. These findings together with our results presented here suggest that the zigzagging exhibited by flying insects, and likely other flying and swimming animals too, is crucial for them to estimate the ambient wind direction in order to guide plume tracking decisions.

### C. Optimal choice of time constants

Our active anemosensing hypothesis involves the assimilation of sensory measurements across three different time constants: *ω, τ*, and *T*. Our results suggest that a good rule of thumb for accurate ambient wind direction estimates is to choose *ω* = turning frequency, and *τ* = *T* ≥ 2/*ω*. This ensures that wind direction estimates are always based on information that spans at least one left or right turn. With noisier sensor measurements, slower turning frequencies and longer values of *τ* are helpful. The increased errors associated with shorter values of *τ* can be partially mitigated by using a much larger value for *T*.

### D. Simplified mathematics with constant velocity control

It is unlikely that insects use the approach we outline in Fig. 3 due to the computational expense (solving the estimation for *τ* = *T* = 10 & *dt* = 0.1 takes ≈ 160 ± 20 milliseconds on a powerful desktop computer). Here we discuss some alternatives. Although plume tracking insects exhibit variability in their ground speed (e.g. Fig. 5E and [44]), it is also well established that they use visual feedback in order to try and maintain approximately constant ground speeds [35], [36], [45]. By assuming both a constant ground speed (*g*) and wind speed (*w*) over the period *τ*, their ratio (*v* = *g/w*) will also be constant. Assuming constant *v* it is possible to dramatically simplify the mathematics, as detailed in the following subsections. Future work will be required to determine how sensitive these approaches are to breaking their assumption of 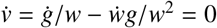.

#### 1) Quadrant sensing

The first simplification we consider does not aim to precisely determine the ambient wind direction, instead it only identifies which 90° quadrant the wind direction lies inside. This requires two comparisons of signs. First, recall the following equation (derived from Eqn. 3 with *δ* = *γ* – *ψ* and *α* = *γ* + *ϕ*):

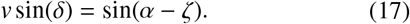

Although *v* is unknown, it is defined as a magnitude and must be > 0, so we can write:

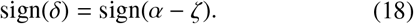

This relationship can be used to determine which hemisphere the ambient wind is coming from, and is the same constraint we use in our convex optimization formulation to disambiguate between up- and downwind. To further constrain the wind direction to one of four quadrants, we can place a similar constraint on the derivative of Eqn. 17 (assuming 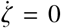 over the period of time over which the derivative is calculated):

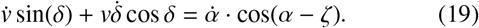

We can now place the same type of sign constraint on this equation:

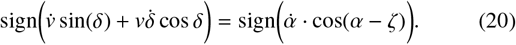

As long as 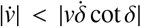, we can ignore the 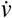 term such i that we have the following constraint (which eliminates the 2 unknown *v* from the equation):

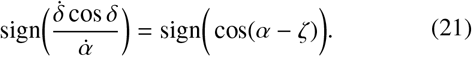

Together, Eqns. 18 and 21 can be used to determine which of four quadrants the ambient wind direction is coming from. Note that although this approach does not provide a precise estimate of wind direction, it may be sufficient for some behaviors, and sequential quadrant estimates could be combined (e.g. through a Bayesian framework) to build a more precise wind direction estimate over time (i.e. over the course of several turns).

#### 2) Discrete least squares solution

Under the assumption of constant *v* = *g/w*, our convex optimization approach (solid orange box, Fig. 3B) reduces to a simple matrix inversion based on Eqn. 14:

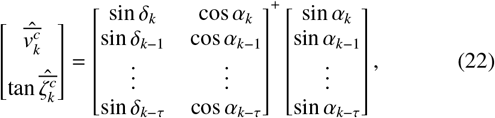

where the ^+^ is the Moore-Penrose pseudoinverse. As with our convex approach (which allows 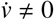), this matrix inversion must be solved for the two cases (tan 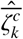and cot 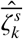), and up- and downwind options must be disambiguated. The quadrant sensing approach could be used in conjunction with least squares estimates of 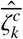 and 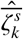 to accurately estimate the wind direction.

Of course, it is unlikely that insects have the neural capacity to invert arbitrary matrices. If we only consider two discrete measurements separated by time *τ* we can write a closed form solution for the 2×2 matrix inversion problem:

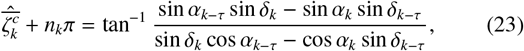

which can be rearranged into the following form to highlight the implicit finite difference calculations that are required,

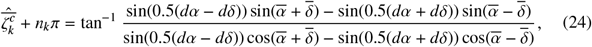

where *dδ* represents a finite difference, *dδ* = *δ_k_* – *δ_k–τ_* and 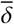 represents a mean calculation, 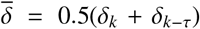, and similarly for *dα*, 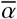.

Because this solution only incorporates measurements from two discrete moments in time, it would likely work best either with filtered measurements to reduce noise, and/or with a filter applied to the resulting 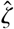 estimates. Furthermore, the best choice of *τ* would likely be one that maximizes the difference between the two measurements as a result of a controlled turn, i.e. *τ* should be some multiple of half the turning period.

#### 3) Continuous direct estimate

A different option (again assuming 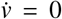) is to recombine Eqn. 9c and Eqn. 10 to define a closed form solution for *ζ* as follows,

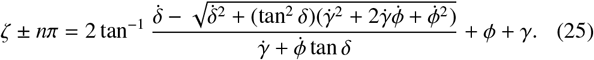

Again, up- and downwind must be disambiguated. This formulation requires sufficiently accurate derivatives of the sensor measurements. As with the quadrant sensing approach, the derivatives could be calculated as a discrete finite difference for measurements taken time *τ* apart, where *τ* is some multiple of half of the turning period.

### E. Considerations for neural implementation

A common theme across each of the simplified estimation approaches outlined above is the presence of a derivative (or finite difference) of sensory measurements. One plausible neural implementation for performing this calculation would be through a mechanism similar to the delay-and-correlate circuit employed for optic flow calculations [46], [47]. Our analysis of different trajectories found that it is important for sensory information to be incorporated over a sufficiently large time constant, *τ*, such that information is assimilated from both sides of at least one turn. Thus, if insects do employ a delay-and-correlate mechanism, we would expect the delay to be either synchronized with voluntary turns, or to be hard-coded as one or multiple values with a time constant equal to half of the typical turning period (i.e. 1 Hz based on Fig. 5).

Each of the three simplified approaches require evaluating complex mathematical expressions containing trigonometric functions as well as multiplication and division. How might the brain perform these computations? In principle, a feedforward neural network can be used to approximate any of these mathematical expressions [48], however, accurate approximations may require too many elements to be practical. Instead, some simplifications or alternative reference frames may be used. The tangent function, for example, could be evaluated as a ratio of two orthogonal vectors, and recent work has revealed how flies can perform certain vector calculations related to navigation [16], [49]. Multiplication and division in general can be converted to addition and subtraction if the calculations are performed in a log-space. Finally, it is quite possible that for the limited range of likely inputs an insect might actually experience due to environmental, biomechanical, or behavioral constraints, the true mathematical expressions could potentially be simplified without introducing excessive errors.

### F. Limitations

In our analysis we only considered kinematically prescribed trajectories. Although flying insects do have exceptional gust rejection abilities [31], they will be thrown off course when subject to strong enough gusts. In such cases, it could be that estimation accuracy of wind direction actually improves due to the change in course direction, though only if the wind is not so overpowering that the insects end up drifting with the wind [23].

We focused our active anemosensing estimation framework exclusively on angular measurements known to be encoded in the central complex. A more traditional estimation approach from engineering would combine control inputs with these available measurements and a detailed dynamics model of the relevant physics to provide more accurate estimates (e.g. with a Kalman filter [50]). It is conceivable that insects could incorporate copies of their motor control commands with a model of the muscle output and aerodynamic forces to more accurately estimate the wind direction, however, this would require detailed calibrations of all of the model parameters. As with the sensory system, these parameters will vary in time due to damage [51], temperature fluctuations [52], or other circumstances, making it challenging to rely on a model-based approach. A significant advantage of relying exclusively on angular measurements is that the entire angular sensorimotor pathway relevant to ambient wind estimation can be self-calibrated without external help [23].

### G. Hypotheses and predictions

If flies exclusively rely on visual anemotaxis, they would not be able to make arbitrary changes in course with respect to the ambient wind direction. Instead, they would need to determine upwind, or perform rhythmic maneuvers centered around the upwind direction. Under our active anemosensing hypothesis, however, flies could continuously adjust their course angle without first orienting upwind.

With respect to computational complexity, the three estimation approaches that assume constant *v* = *g/w* provide the most biologically plausible implementations of our active anemosensing hypothesis. In each of these cases we would expect to find a clear encoding of *δ* = *γ* – *ψ* in the flies’ brain. Furthermore, we would expect to find neural pathways capable of differentiating this signal with a time constant roughly equal to half their stereotypical turning period of ~ 2 sec.

We found that flies typically cast at approximately 0.5 Hz, i.e. ~ 1 left and ~ 1 right turn every two seconds. Our active anemosensing hypothesis suggests that for this turning frequency, flies would have most accurate wind direction estimates for a *τ* ≥ 1 sec. This would allow flies to be effective at tracking plumes in highly dynamic wind that changes direction on the scale of 1 second, possibly explaining the release and recapture successes at low wind speeds described previously. For insects with noisier sensory measurements, either due to fundamental sensor limits or environmental constraints such as low light levels, we predict that the turning frequency should decrease. Larger plume tracking animals such as albatross [53] could also conceivably employ active anemosensing to estimate ambient wind direction. With slower dynamics due to their more massive bodies we would also expect to see slower turning frequencies. For both cases, we predict that slower turning frequencies should lead to larger required values of *τ* and thus slower responses to rapid changes in wind direction.

One particular challenge our analysis highlights is that of distinguishing up- vs. downwind. The most challenging scenario is for an insect to distinguish between flying downwind faster than the wind speed, vs. flying upwind. To resolve this challenge, insects could use a similar approach to our solution (Eqn. 15), or perhaps they employ a combination of active anemosensing and visual anemotaxis to guarantee that during plume tracking behaviors they are always generally headed upwind, this would be equivalent to using a very large value of *T*.

## Acknowledgements

We thank Ethan Swierski for help with developing and testing the drone, and Katherine Nagel for helpful discussions and feedback on the manuscript.

## VII. Data accessibility statement

All of the code associated with this paper is provided in an open source repository here: https://github.com/florisvb/active_anemosensing [54]. Data is available as a Data Dryad repository here: https://doi.org/10.5061/dryad.gb5mkkwrv, and will be available upon publication. For review, temporary data access is available at: https://datadryad.org/stash/share/GJm64N6jRFiD6j43ewAhM18XMTJbpyvmP3uCV-PhOaY.

## VIII. Supplemental Materials

The following two subsections provide a more in depth mathematical analysis of the visual anemotaxis hypothesis.

### A. Visual anemotaxis phase portraits in 2-dimensions

Here we provide a more complete analysis of the dynamics of visual anemotaxis for different gain constants. From Eqn. 3 and Eqn. 5 we construct the following relationship,

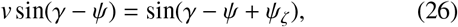

where *ψ_ζ_* is the global course direction relative to the wind direction (Eqn. 5). Combining Eqn. 4, and Eqn. 6 we can construct the following relationship,

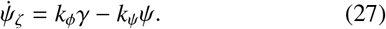

Solving for *γ* yields:

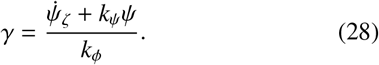

Plugging Eqn. 28 into Eqn. 26 and solving for 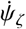 yields:

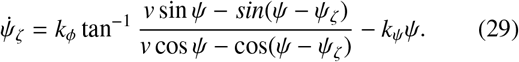

Together with Eqn. 4B 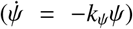 we now have a 2-degree of freedom system (the free variables being *ψ* & *ψ_ζ_*). Figure S1 shows the 2-dimensional phase portrait for this system given different choices of *v, k_ψ_, k_ϕ_*. So long as the gain constants are greater than zero, the system has stable fixed points corresponding to movement upwind, and unstable fixed points corresponding to movement downwind. When the two gain constants are different from one another the trajectories in phase space become less direct, and when *v* (the ratio of ground speed and wind speed) is large the trajectories in phase space are also less direct.

### B. Visual anemotaxis with an offset from upwind

In order to use the visual anemotaxis strategy to move in a direction offset from upwind, insects could incorporate an angular offset, *ϵ*, into the control of *ϕ*:

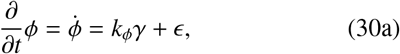

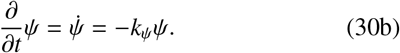

Repeating the analysis that led to Eqn. 8 we arrive at:

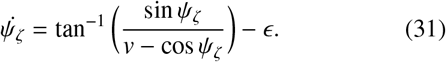

Figure S2 shows the 1-dimensional phase portrait for this relationship for different choices of *ϵ*. In this case, the stable fixed points are offset from upwind, however, the offset is a function of both *ϵ* and *v*. Thus, although this strategy would allow an insect to maintain flight at an angle relative to the ambient wind, the insect would not have access to what this angle is by using visual anemotaxis alone. In order for an insect to fly at a specific desired angle relative to upwind using visual anemotaxis, it would first need to orient upwind, and then change course while remembering the global wind direction.

## IX. Supplemental figures

**Fig. S1:**
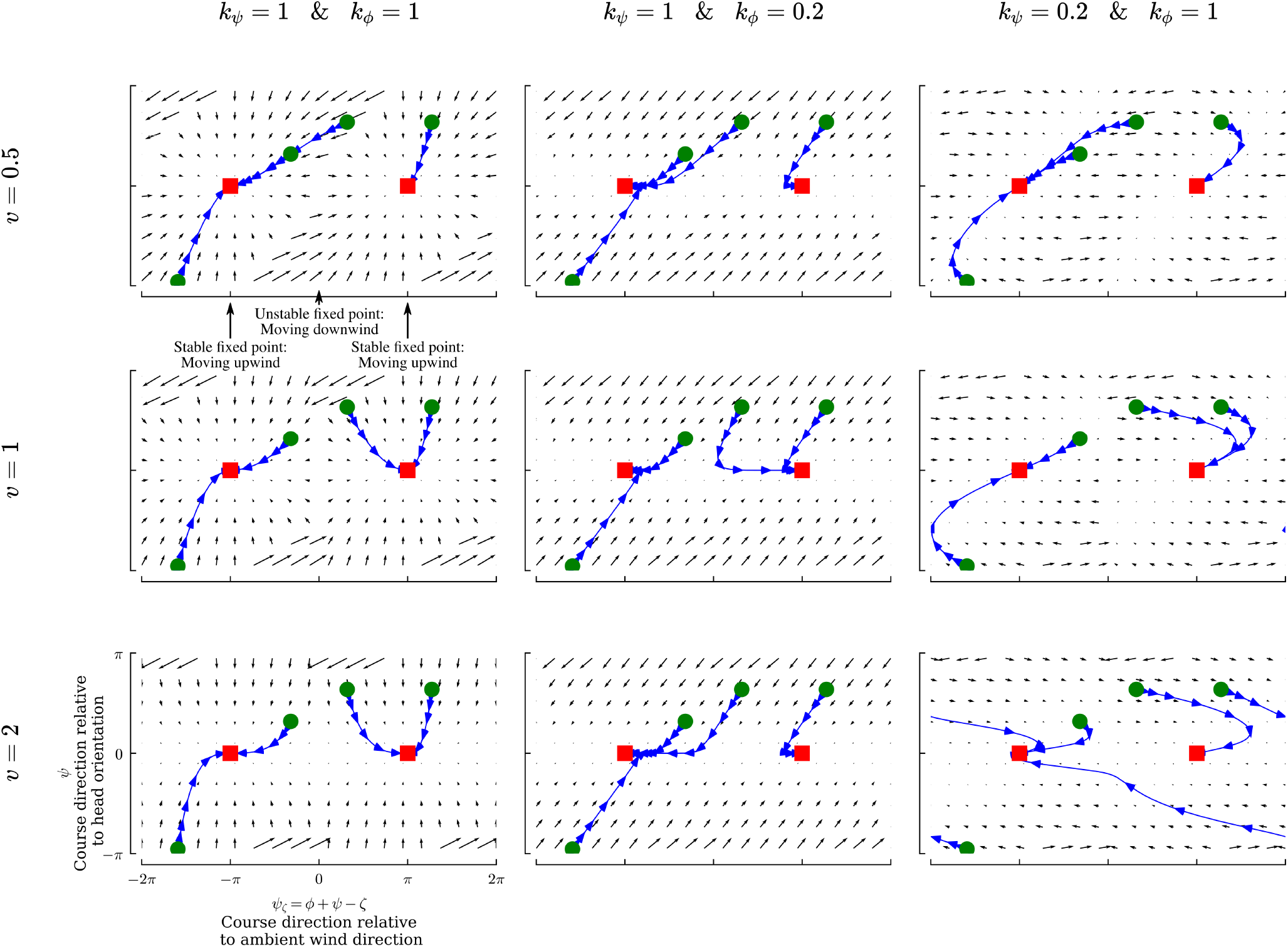
2-dimensional phase portraits illustrating the stability of the visual anemotaxis algorithm for different gain constants (*k_ψ,ϕ_*) and different non-dimensional velocities (*v* = *g/w*). Each panel shows the 2-dimensional phase portrait (black arrows) for the direction of motion relative to the head orientation (*ψ*) and the global direction of motion relative to the wind direction (*ψ_ζ_*). Blue lines show the flow of 4 trajectories along with their initial condition (green circle) and final position (red square).

**Fig. S2:**
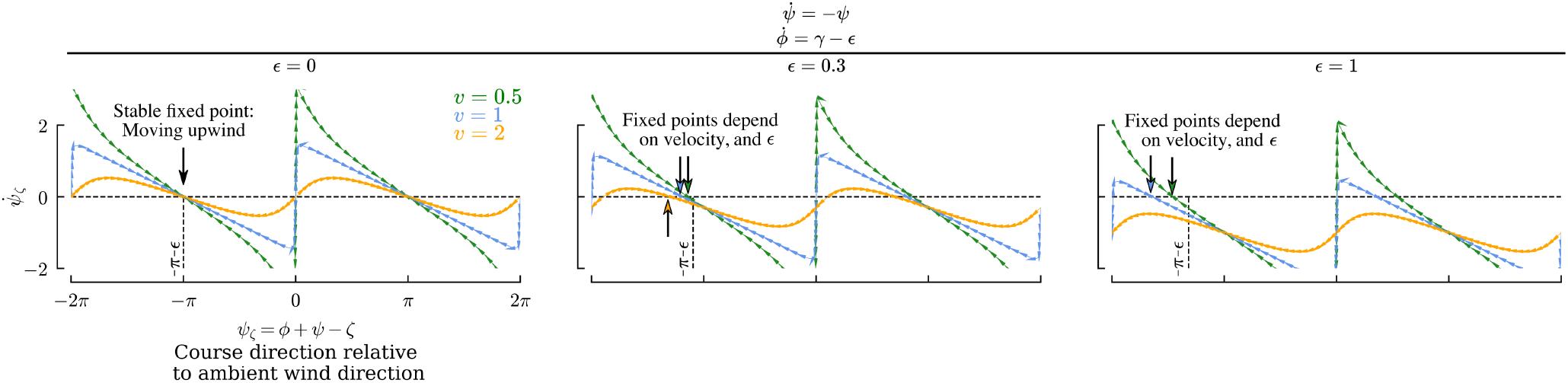
1-dimensional phase portraits, plotted as in Figure 2, for different offsets (*ϵ*) in the control of the orientation *ϕ*. Note how the fixed points depend on both *v* and *ϵ*, and that in some cases for larger values of *v* there are no stable or unstable equilibria.

**Fig. S3:**
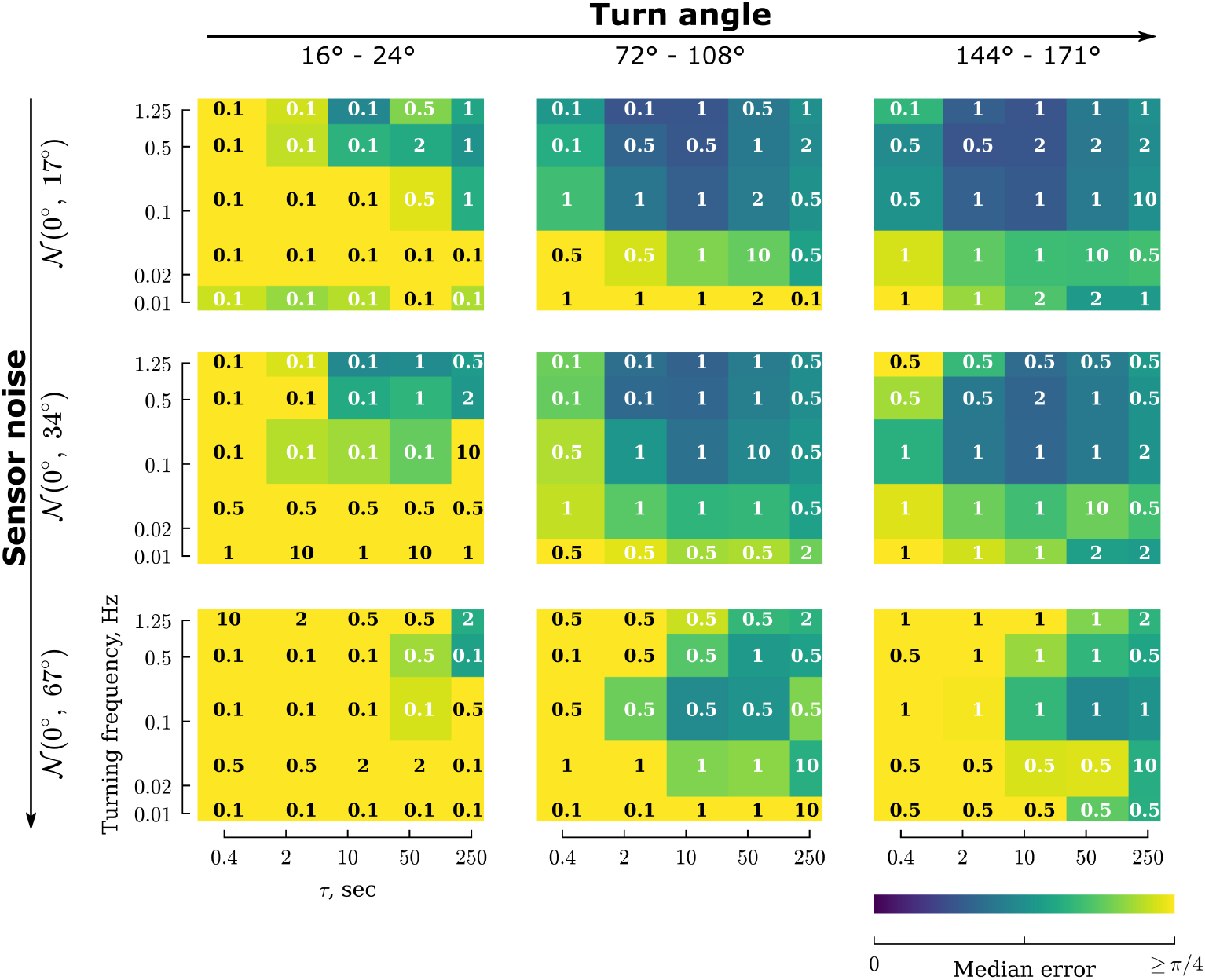
Optimal choice of turning frequency and *τ* are minimally effected by the precise choice of *ω*. Heatmaps are plotted as in Fig. 8B for three types of trajectories and noise levels. Comparing this figure with Fig. 9 shows that the precise choice of *ω* has little effect on the median error.

**Fig. S4:**
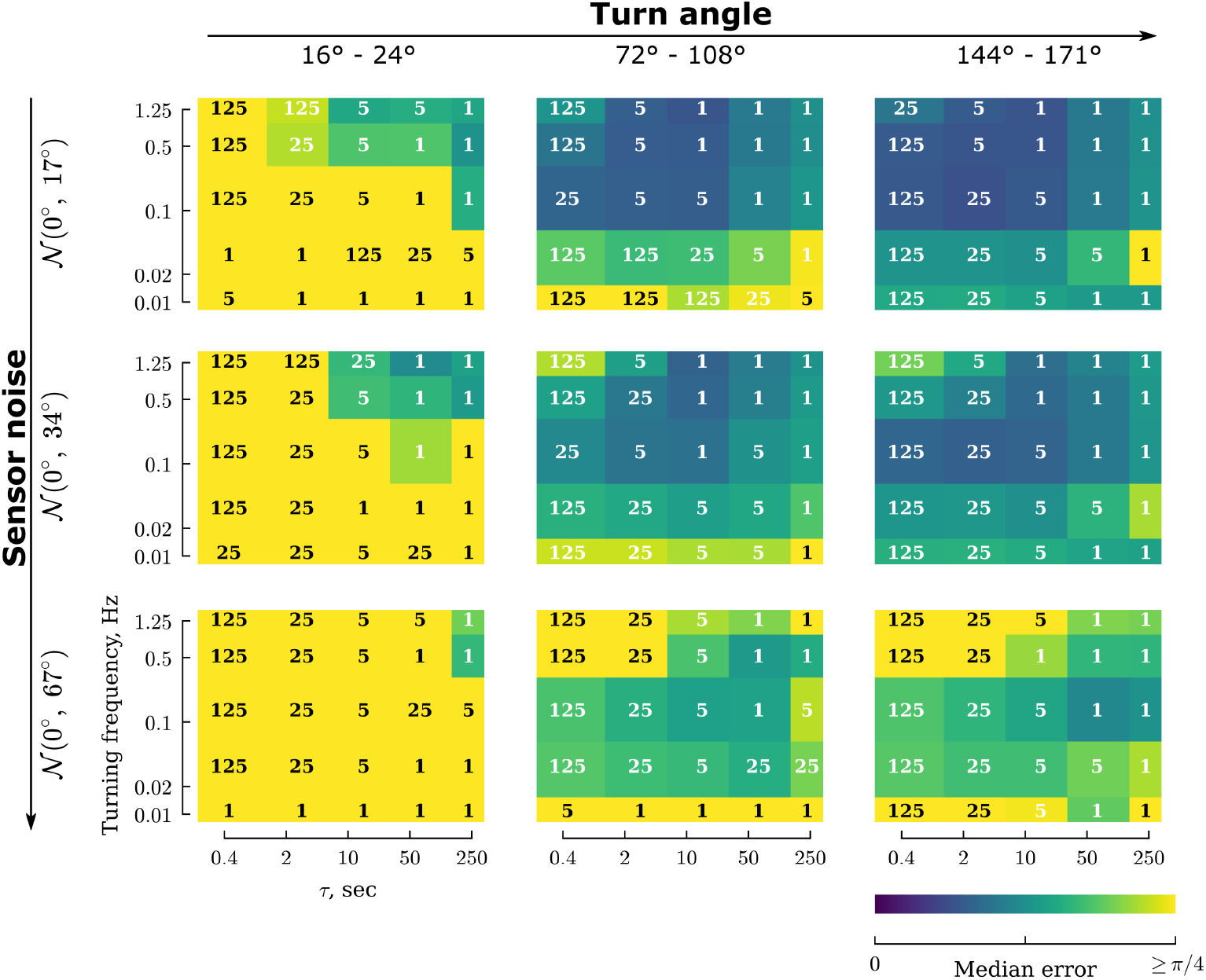
For trajectories with medium and large turn angles, ambient wind direction estimates can be improved for small *τ* scenarios by using a much larger value of *T* to disambiguate between up- and down-wind. Heatmaps are plotted as in Fig. 8C for three types of trajectories and noise levels.

**Fig. S5:**
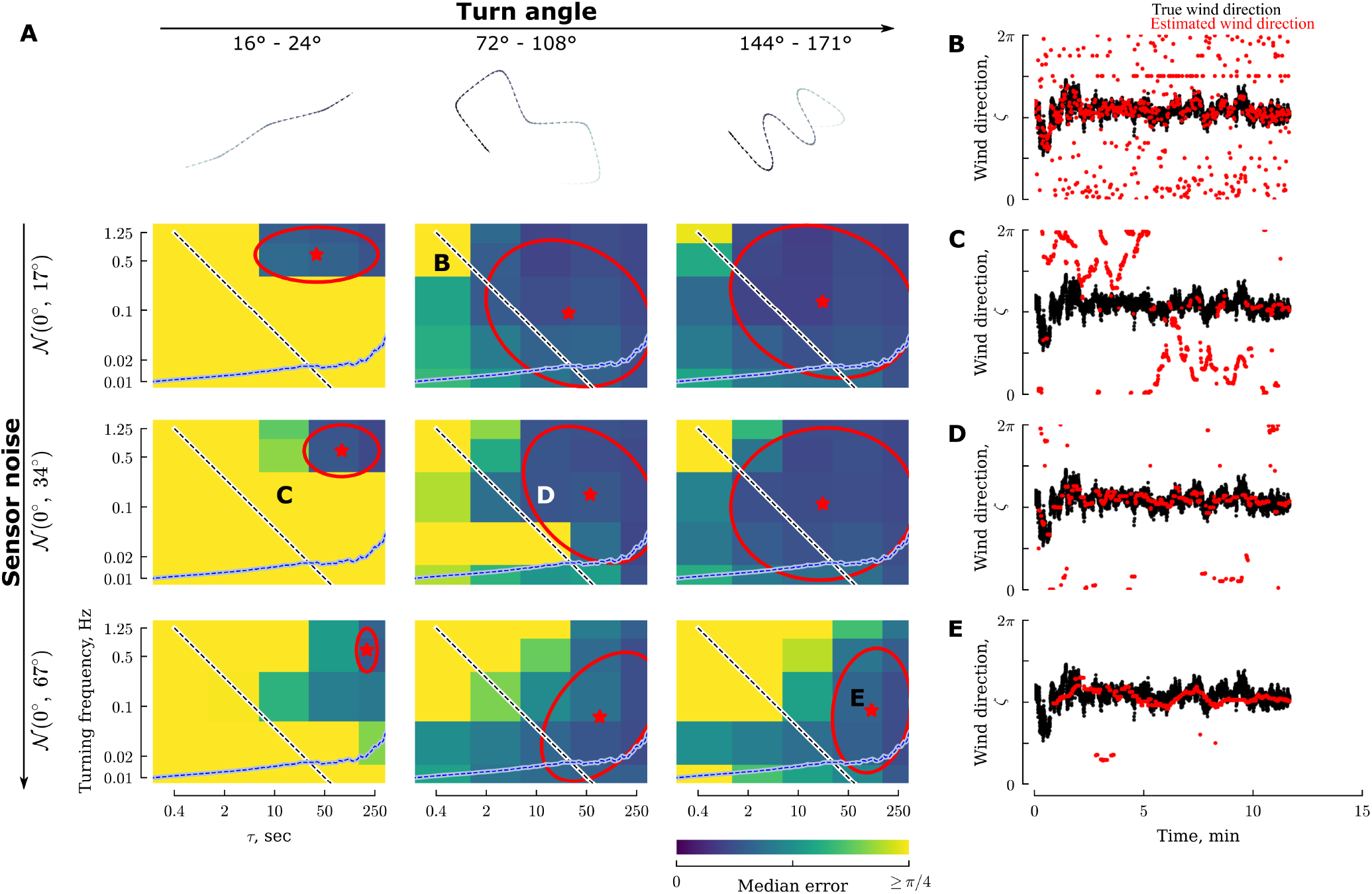
In consistent wind scenarios, errors in ambient wind direction estimates are lower, larger values of *τ* are permissable, as are slower turning frequencies. Data plotted as in Fig. 9

**Fig. S6:**
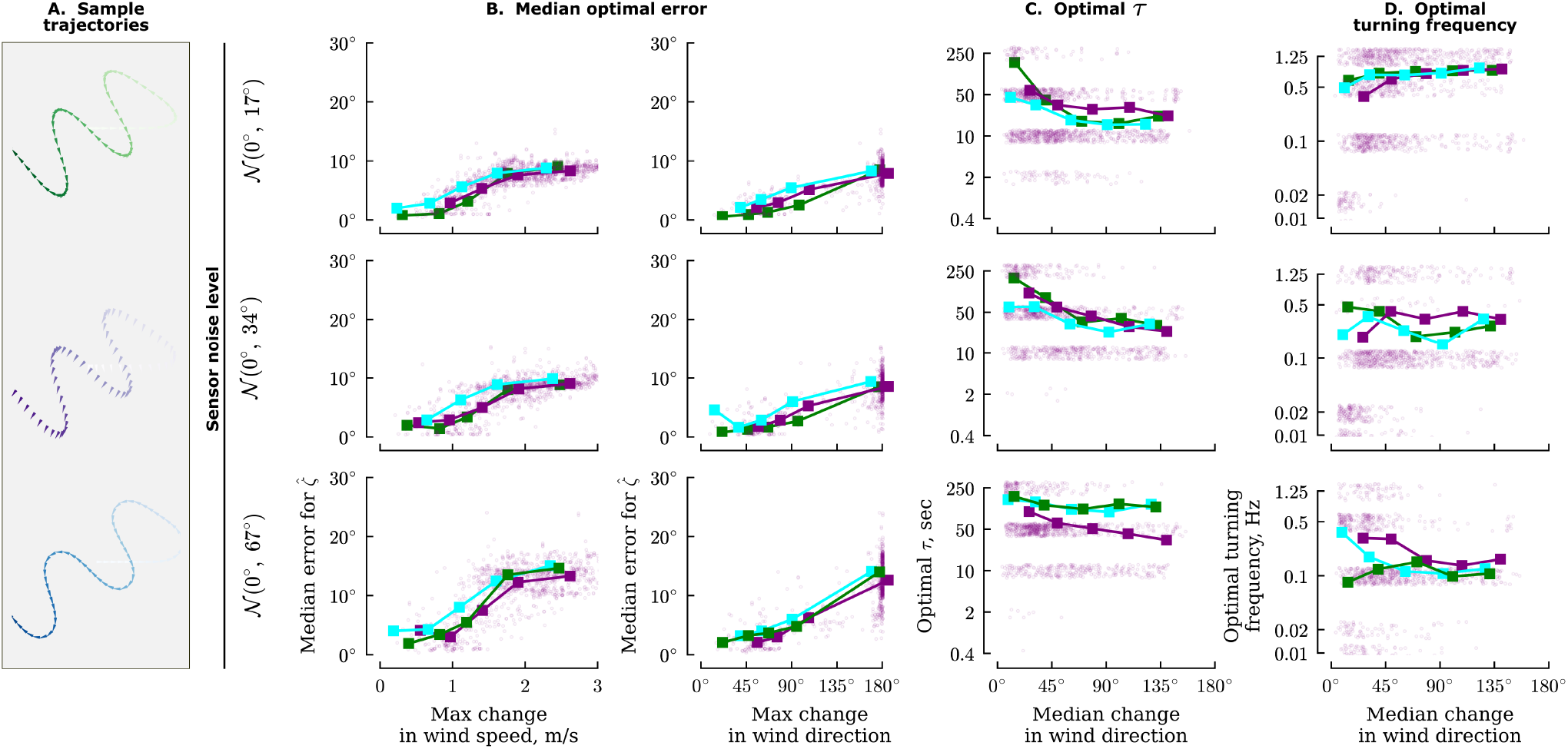
Summary of optimal error, *τ*, and turning frequency as a function of sensor noise, and wind variability for different trajectory types. Results plotted as in Fig. 11. The green colored data are repeated from Fig. 11, and are provided for reference.

**Fig. S7:**
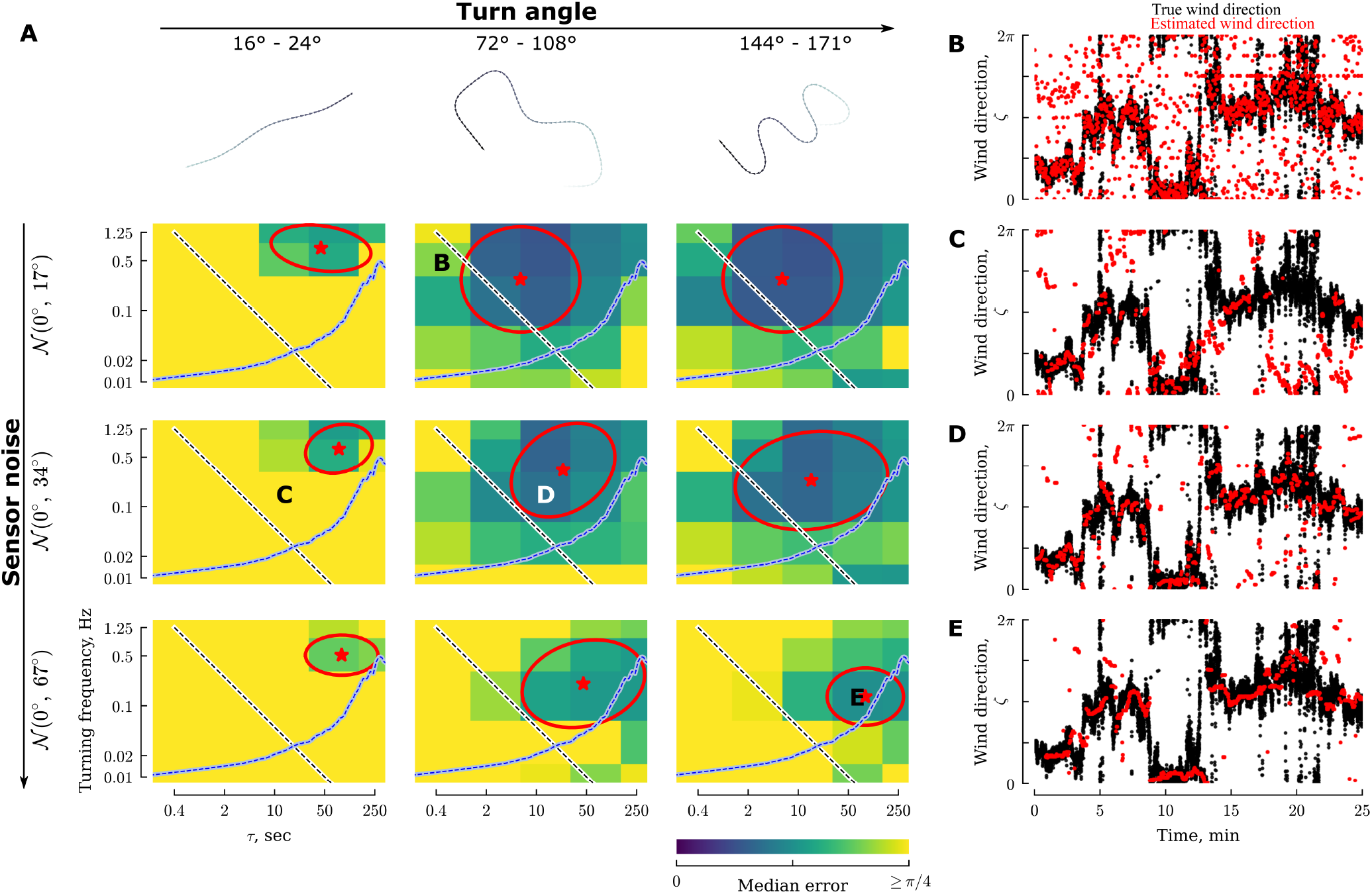
Constant velocity trajectories do not offer any advantage in estimation accuracy compared to variable velocity trajectories. Data plotted as in Fig. 9

**Fig. S8:**
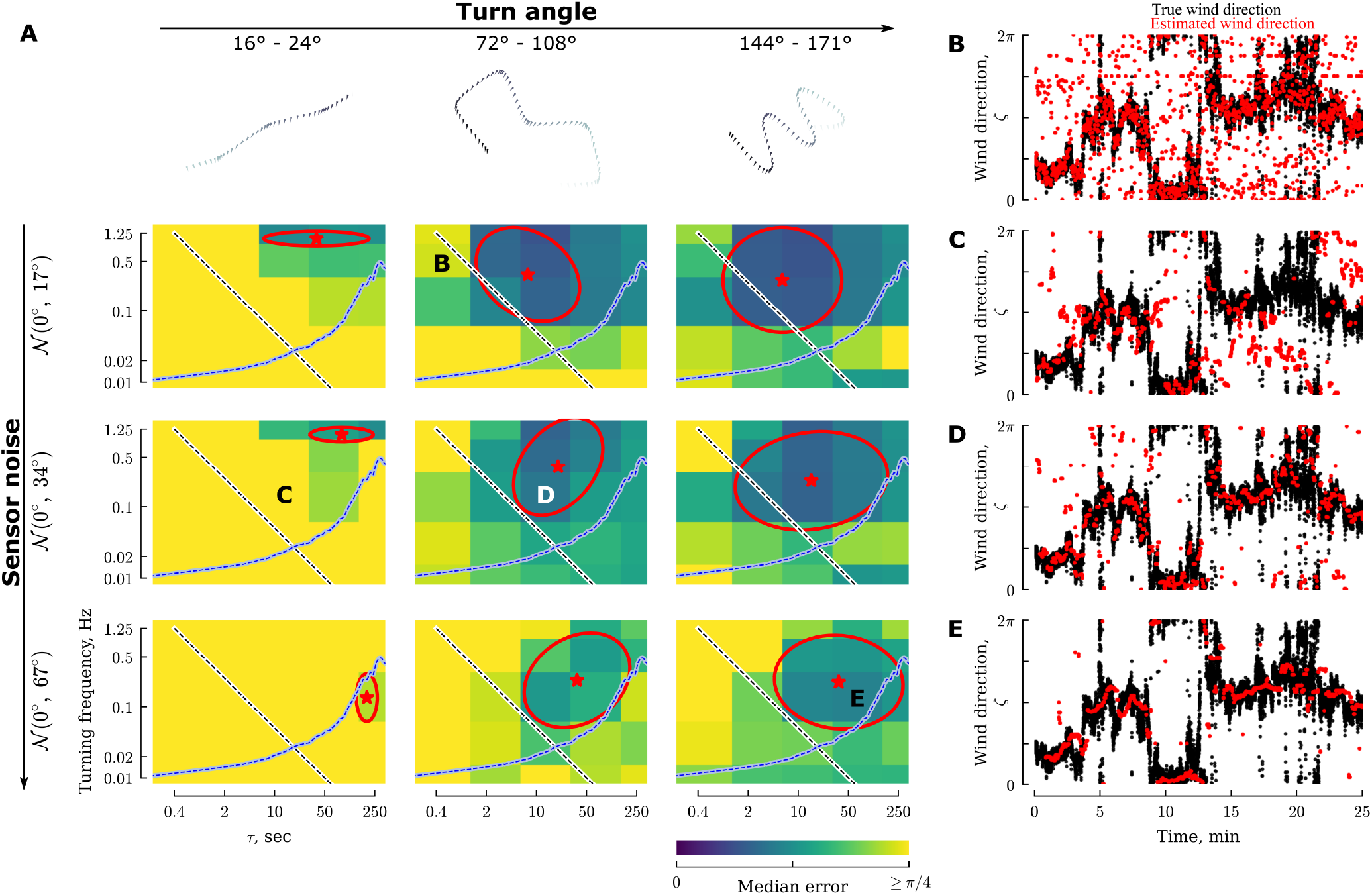
Constant *γ* trajectories offer a subtle improvement in estimation accuracy for scenarios with large turns and high noise. Specifically, lower estimation errors can be achieved for smaller values of *τ*. See text for detailed discussion. Data plotted as in Fig. 9

**Fig. S9:**
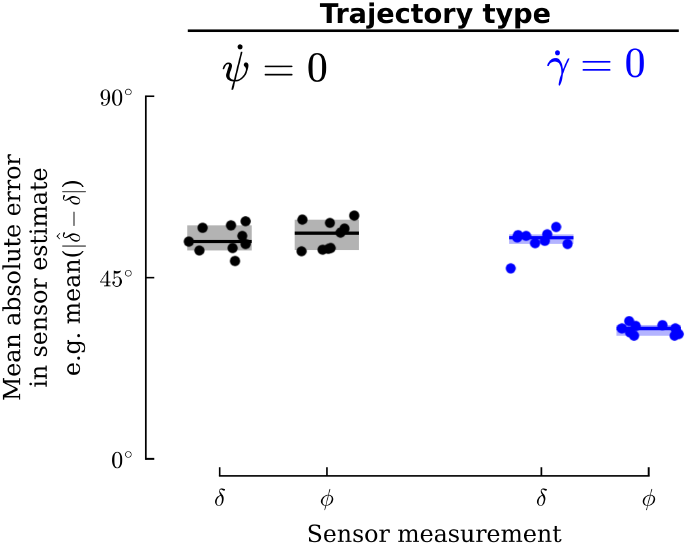
Mean absolute error in estimates of *δ* = *γ* – *ψ* and *ϕ* for constant-*ψ* and constant-*γ* trajectories with large turn angles (~ 160°), a turning frequency of 0.1 Hz, and Gaussian noise with a standard deviation of 67°, for the dynamic wind scenario, for 10 different random seeds. Lines and shading show the median and 95% confidence interval. The absolute errors of the smoothed estimates (*ω* = 0.1 Hz) for *δ* are similar for the two cases, but the smoothed estimates for *ϕ* have half the error for the constant *γ* trajectories.

